# Advancing knowledge of West African morning glories: a taxonomic revision of *Ipomoea* L. (Convolvulaceae) from Ghana

**DOI:** 10.1101/2024.04.02.587553

**Authors:** Benjamin Darko Williams, Ruth Cristóvão Francisco, Befkadu Mewded, Christiana Peprah Oppong, Claudia Bosomtwi-Ayensu, Collins Wafula Masinde, Deborah Moradeke Chukwuma, Deresa Abetu Gadisa, Doreen Donkor Yeboah, Enangnon Oscar Doré Ahossou, Fitiavana Rasaminirina, Igho-Osagie Uhunwa Precious, Mercy Jebiwott Korir, Kwasi Boateng Antwi-Boasiako, Raymond Attah Mfodwo, Mutegeki Alislam Said Musa, Peter Atta-Adjei, Peter Kwasi Akomatey, Seyiram Kumordzie, Renata Borosova, Cheung Tang, Alex Asase, Gabriel Ameka, Ana Rita Giraldes Simões

## Abstract

Ghana’s plant diversity is estimated at 2,974 plant species, belonging to 1,077 genera and 173 plant families. However, a Flora of Ghana is yet inexistent: targeted floristic and taxonomic studies are still much needed to document the plant diversity of the country fully at the family, generic and species levels. This is essential for identifying priority conservation areas in the country and support further research in crop wild relatives or medicinal plants, which will help tackle food insecurity and improve livelihoods. In this study, we provide a taxonomic revision of the *Ipomoea spp.* in Ghana to enhance their identification, conservation and sustainable utilization as food and medicine among other uses. An extensive literature review was carried out, including historical references and online taxonomic databases, to recover information on accepted names, type specimens and synonyms, followed by consultation of herbarium specimens at GC herbaria, to retrieve morphological information and database specimens. Specimen locality information was georeferenced, and records plotted onto distribution maps. As a result, this work provides an identification key to the species of *Ipomoea* of Ghana, nomenclatural information, comprehensive morphological descriptions, detailed list of examined specimens, distribution maps and notes on conservation status and traditional plant uses. In total, 28 species are fully described, 20 of which are native and eight introduced from the Americas; five are new records to Ghana.

## Introduction

Ghana is located in West Africa, bounded by Burkina Faso on the north, Togo on the eastern border, Côte d’Ivoire on the west, and the Gulf of Guinea on the south. It is home to 2,974 indigenous species of flowering plants, belonging to 1,077 genera and 173 families (Ministry of Environment, Science and Education, 2016). From south to north, three main vegetation types can be found: forest zone, the northern savanna zone, and the coastal savanna (Asase *et al*., 2010). There are also many types of biomes, but the tropical high forests and the savannas are the two major areas represented (Ministry of Environment and Science, 2002). Ghana has a total surface area of 238,540 km^2^; half of the country is below 152m (499 feet) above sea level, and the highest point is 883m above sea level; the average monthly rainfall in June ranges from 152 to 254mm; January is a dry month, while August is the driest month in the eastern coastal regions (Ghana Meteorological Agency, 2022).

Convolvulaceae, commonly known as the plant family of “morning glories’’ and “sweet potato” is an important source of food, medicine and several ornamental plants. It comprises ca. 1,977 species distributed among 60 genera (Eserman *et al*., 2020), occurring across tropical and temperate regions. They can be herbs, shrubs, or twiners with alternate leaves, actinomorphic (4) 5 – merous flower; usually free and overlapping sepals, tubular corolla with distinct midpetaline bands, superior ovary, and usually dehiscent 4-seeded capsular fruit (Staples & Brummitt, 2007; Bramley, 2020; Simões 2023).

*Ipomoea* L. is the largest genus in the family Convolvulaceae with 635 accepted species; the species of *Ipomoea* are ubiquitous, spreading majorly in tropical and subtropical regions and some concentrated in temperate regions (Eserman *et al*., 2020; Wood *et al.,* 2020; POWO, 2024). Economically, *Ipomoea* is of intrinsic economical value ranging from ornamentals, medicinal to food plants (Gill & Nyawuame, 1991; Folorunso, 2013; Sultana & Rahman, 2016; Wood *et al*., 2020). *Ipomoea purpurea* (L.) Roth, *Ipomoea batatas* (L.) Lam. and *Ipomoea violacea* L., for example, are exploited for their ornamentals, culinary, religious ritual and medicinal values (Eserman *et al*., 2020; Srivastava & Rauniyar, 2020; Wood *et al*., 2020). Morphologically, species in *Ipomoea* are mostly herbaceous to woody twiners or small trees, with great variation in corolla shape and colour, as well as other vegetative and reproductive characters (Meira *et al*. 2012; Eserman *et al.,* 2020). It is also the genus of “sweet potato” (*Ipomoea batatas* (L.) Lam.), an important crop and important nutritional source all over the world (Joseph & Anthony, 2014). In many African countries, like Ghana, they serve as a highly nutritious source of food that has been used to curb hunger and malnutrition (Darkwa & Darkwa, 2013); in East Africa, the demand for sweet potato has been increasing with the growing interest in “super vegetables’’ (Cernansky, 2015).

However, many taxonomic and conservation gaps exist in this genus, especially in tropical Africa. A recent monographic work of *Ipomoea* focused mostly on the American continent (Wood *et al.,* 2020), excluding the greatest part of African and Asian species. Thus, of the 635 species of *Ipomoea* currently accepted (POWO, 2024), only 72 have had their conservation status assessed, which corresponds to c. 11% of the accepted species. Of those assessed, six are Critically Endangered, 15 Endangered, and nine Vulnerable (IUCN, accessed February 2024). Therefore, 89% of the species of *Ipomoea* across its global distribution remain of unknown conservation status.

For West Africa, the latest available flora account of Convolvulaceae is the Flora of West Tropical Africa, reporting 75 species belonging to 16 genera; of these, 41 species (more than half) belonging in genus *Ipomoea* (Heine, 1963). However, six decades later, the available literature for Convolvulaceae in West Tropical Africa is yet scanty, with no comprehensive taxonomic revisions for the family or of *Ipomoea* species in the region so far. For Ghana, there is not yet a Flora account of the flowering plants of the country, and neither a taxonomic treatment of Convolvulaceae. Hence, the number of species, where they occur, their economic or traditional use, and their conservation status is not yet fully documented, undermining broader studies on the floristic diversity of Ghana, conservation efforts and further research on its useful plants.

The overall goal of this research work is to compile detailed and reliable information on species of *Ipomoea* L. in Ghana, a diverse and widely economically important genus that includes morning glories and sweet potato wild relatives, to enhance their identification, conservation and sustainable utilization as food and medicine. An identification key, full morphological descriptions, and notes on distribution, conservation and traditional uses are provided for all species of *Ipomoea* L. that we were able to document in Ghana through a variety of sources: taxonomic literature, analysis of herbarium collections at GC herbarium in Accra, Ghana and taxonomic online resources.

*Ipomoea* is represented by 28 species in Ghana: of these, 20 are native and eight introduced from the Americas, presumably for their ornamental, food or medicinal uses (Table I). Five are recorded for the first time to occur in Ghana, in respect to existing literature and taxonomic databases, two of which are native (*I. pyrophila* A. Chev. And *I. stenobasis* Brenan) and three presumed introduced from the Americas (*I. intrapilosa* Rose, *I. quamoclit* L. and *I. triloba* L.). All species are considered Least Concern, except for *Ipomoea batatas* (sweet potato) which has been previously assessed as Data Deficient (Rowe *et al.,* 2019).

**Table I.**
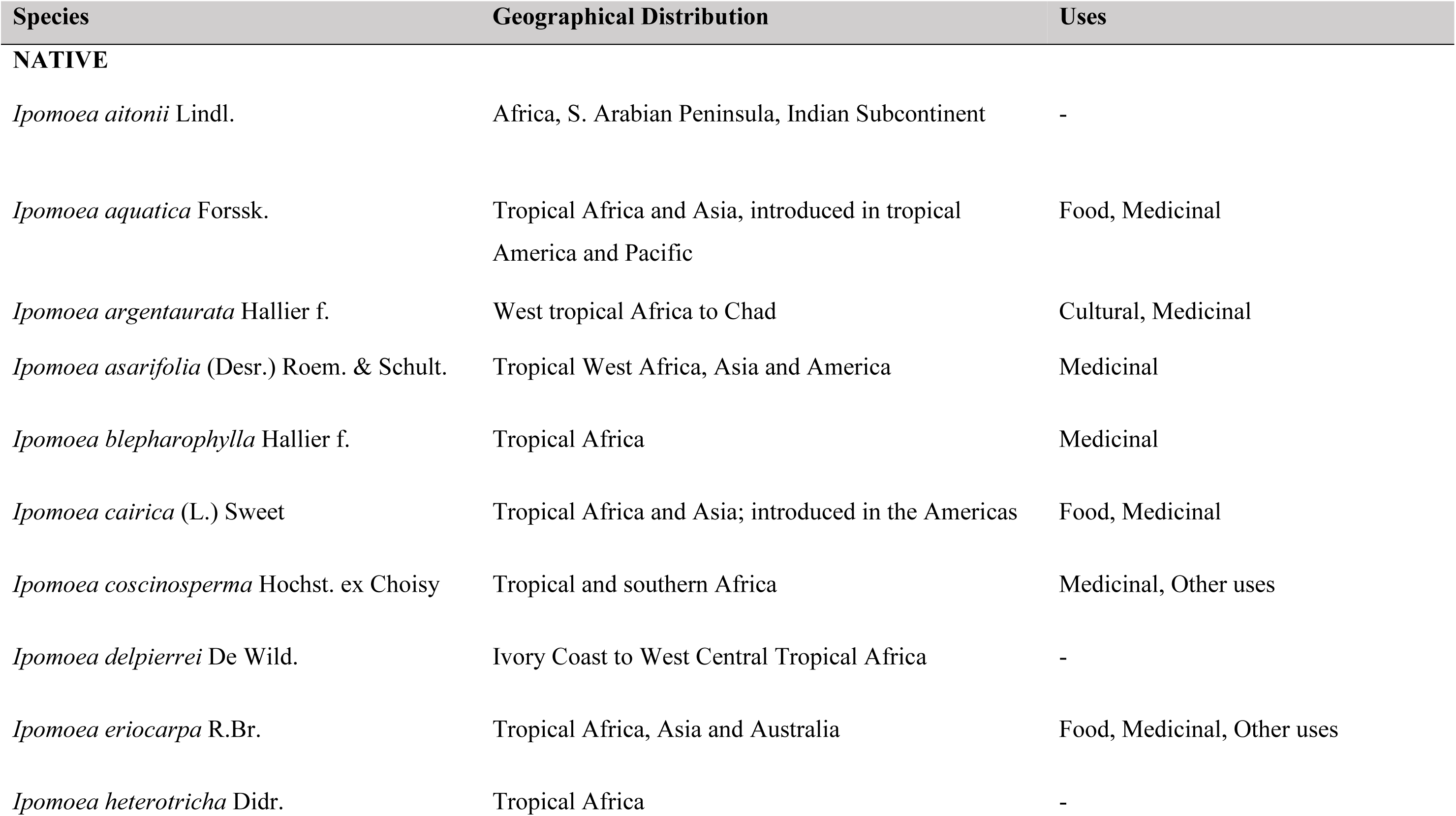
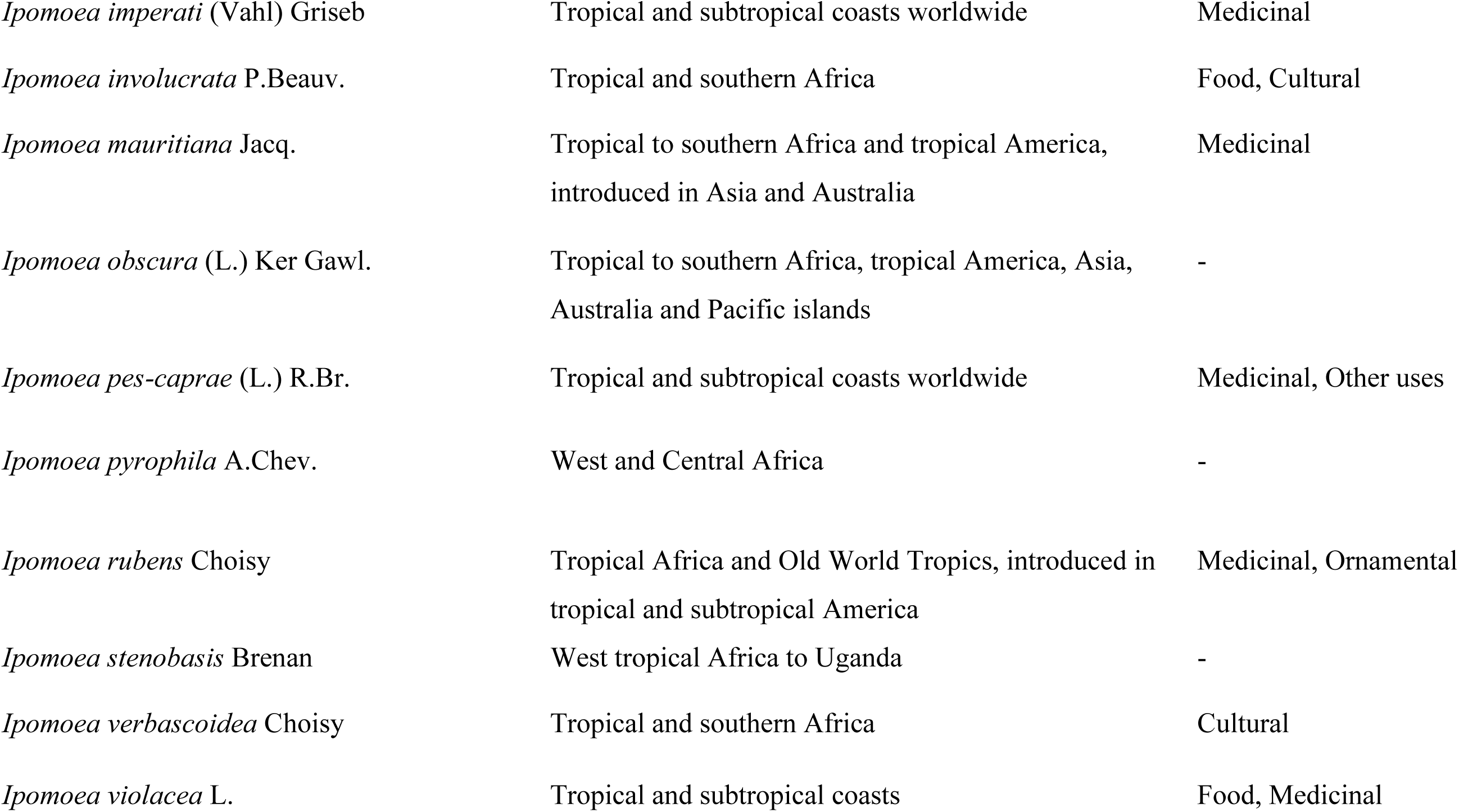
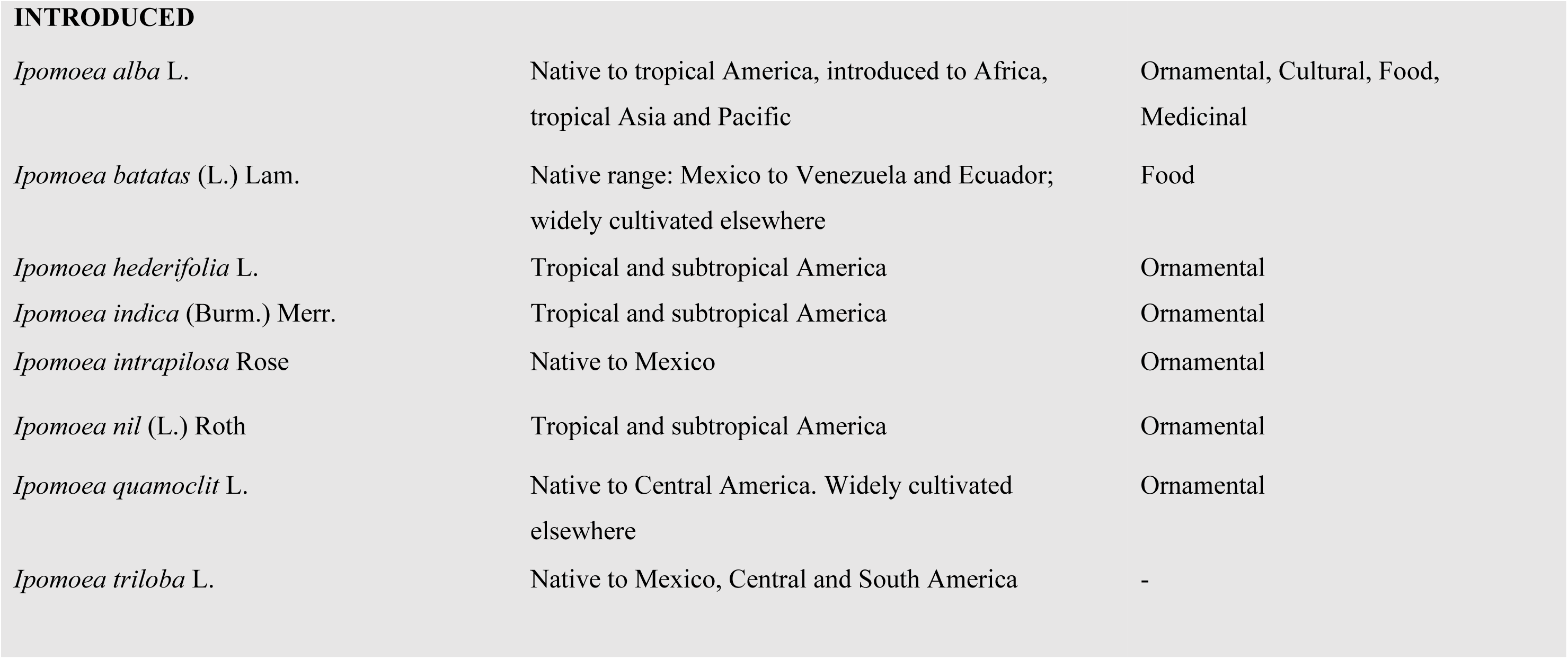
List of species of *Ipomoea* in Ghana, their distribution and known uses.

| Species | Geographical Distribution | Uses |
| --- | --- | --- |
| **NATIVE** |  |  |
| *Ipomoea aitonii* Lindl. | Africa, S. Arabian Peninsula, Indian Subcontinent | - |
| *Ipomoea aquatica* Forssk. | Tropical Africa and Asia, introduced in tropical America and Pacific | Food, Medicinal |
| *Ipomoea argenteaurata* Hallier f. | West tropical Africa to Chad | Cultural, Medicinal |
| *Ipomoea asarifolia* (Desr.) Roem. & Schult. | Tropical West Africa, Asia and America | Medicinal |
| *Ipomoea blepharophylla* Hallier f. | Tropical Africa | Medicinal |
| *Ipomoea cairica* (L.) Sweet | Tropical Africa and Asia; introduced in the Americas | Food, Medicinal |
| *Ipomoea coscinosperma* Hochst. ex Choisy | Tropical and southern Africa | Medicinal, Other uses |
| *Ipomoea delpierrei* De Wild. | Ivory Coast to West Central Tropical Africa | - |
| *Ipomoea eriocarpa* R.Br. | Tropical Africa, Asia and Australia | Food, Medicinal, Other uses |
| *Ipomoea heterotricha* Didr. | Tropical Africa | - |
| *Ipomoea imperati* (Vahl) Griseb | Tropical and subtropical coasts worldwide | Medicinal |
| *Ipomoea involucrata* P.Beauv. | Tropical and southern Africa | Food, Cultural |
| *Ipomoea mauritiana* Jacq. | Tropical to southern Africa and tropical America, introduced in Asia and Australia | Medicinal |
| *Ipomoea obscura* (L.) Ker Gawl. | Tropical to southern Africa, tropical America, Asia, Australia and Pacific islands | - |
| *Ipomoea pes-caprae* (L.) R.Br. | Tropical and subtropical coasts worldwide | Medicinal, Other uses |
| *Ipomoea pyrophila* A.Chev. | West and Central Africa | - |
| *Ipomoea rubens* Choisy | Tropical Africa and Old World Tropics, introduced in tropical and subtropical America | Medicinal, Ornamental |
| *Ipomoea stenobasis* Brenan | West tropical Africa to Uganda | - |
| *Ipomoea verbascoidea* Choisy | Tropical and southern Africa | Cultural |
| *Ipomoea violacea* L. | Tropical and subtropical coasts | Food, Medicinal |
| INTRODUCED |  |  |
| *Ipomoea alba* L. | Native to tropical America, introduced to Africa, tropical Asia and Pacific | Ornamental, Cultural, Food, Medicinal |
| *Ipomoea batatas* (L.) Lam. | Native range: Mexico to Venezuela and Ecuador; widely cultivated elsewhere | Food |
| *Ipomoea hederifolia* L. | Tropical and subtropical America | Ornamental |
| *Ipomoea indica* (Burm.) Merr. | Tropical and subtropical America | Ornamental |
| *Ipomoea intrapilosa* Rose | Native to Mexico | Ornamental |
| *Ipomoea nil* (L.) Roth | Tropical and subtropical America | Ornamental |
| *Ipomoea quamoclit* L. | Native to Central America. Widely cultivated elsewhere | Ornamental |
| *Ipomoea triloba* L. | Native to Mexico, Central and South America | - |

## Material and Methods

The present taxonomic work is the result of a group project that was carried out during the “Plant Taxonomy Skills for Ecology and Conservation” course that took place at GC Herbarium in August 2022, jointly organized by Royal Botanic Gardens, Kew and the University of Ghana, with 15 participants from eight African countries (Angola, Benin, Ethiopia, Ghana, Kenya, Nigeria, Uganda and Madagascar). In a first instance, individual participants produced taxonomic treatments for single species of *Ipomoea*, selected among those reported to occur in Ghana in the literature and online databases; later, in groups of five participants, they compiled treatments of five species per group, based on in-person analyses of herbarium collections at GC herbarium.

As a first step for the taxonomic assignment, an extensive literature review was conducted for all species of genus *Ipomoea* L. reported to occur in Ghana, based on taxonomic databases (IPNI, POWO, BHL, JSTOR Global Plants and African Plant Database), floristic treatments from other African regions such as Flora of West Tropical Africa (Heine 1963), Flora of Central Africa (Mwanga-Mwanga, Sosef & Simões, 2022), Flora of Madagascar (Deroin, 2001), Flora of Ethiopia and Eritrea (Demissew, 2006), recent taxonomic works of the genus *Ipomoea* (e.g. Wood *et al.,* 2020) and a wide range of other relevant literature with information on plant uses, chemistry, distribution or conservation status. Thus, available online taxonomic resource tools were also used to confer accepted species names, identify synonyms and locate type specimens (e.g. JSTOR; POWO, 2024,).

Preliminary morphological descriptions were drafted based on the specialised taxonomic literature, and a morphological data matrix was compiled in Microsoft Excel, containing information on morphological characters for each species. A total of 369 plant specimens were consulted from Ghana (GC), which allowed us to revise the morphological data matrix in accordance with the variation found among Ghanaian collections. It also allowed us to identify potential species records not present in the literature. The specimen identification was first confirmed using Floras and other taxonomic literature, and morphological observations were carried out using stereomicroscope, hand lens and ruler. Key morphological characters, like leaf shape, base, margin, apex, size and indumentum, were recorded. The complete morphological descriptions for each of the species were assembled from the morphological data matrix with the aid of a semi-automatic tool (Mail Merge, in Microsoft Word). The species identification key was prepared by comparison of the morphological differences and similarities of the species on the morphological data matrix, and the key was tested against herbarium specimens to ascertain its use for identification.

Information was collected from the labels of all specimens examined, and databased to obtain ancillary information for the taxonomic treatment of each species. The geographical data was retrieved from examined specimens; for specimens that had sparse information on the geographical location, these were further investigated using online tools such as Google Earth (https://www.google.com/earth), with coordinates estimated to the nearest town. Morphological terminology followed Kew Plant Glossary (Bentje, 2010) and herbarium citations followed Index Herbariorum (Thiers, continuously updated).

After the course, the information from all project groups was compiled by adding missing species, updating information against more recent versions of taxonomic databases, and re-analysing GC collections. Then, all authors collaborated remotely in the writing of the manuscript, and refining taxonomic and nomenclatural details.

This work presents the outcome of this course project, although a more comprehensive taxonomic treatment is currently in preparation by the same authors, with extended analysis of additional herbarium specimens from other herbaria, distribution maps and more detailed conservation assessments.

## Taxonomic Treatment

### **Ipomoea** L. (Linnaeus 1753: 159); Linnaeus (1754: 76)

> Type: *Ipomoea pes-tigridis* L.

Annual or perennial *herbs,* or rarely *shrubs*. *Stems* erect, creeping or twining, sometimes lianescent. *Leaves* alternate, simple, rarely compound; leaf blades often cordate at the base, margin entire, lobed or ± deeply divided. *Inflorescence* axillary, in 1-many flowered cymes, umbelliform, corymbiform, paniculiform or in capituliform heads, sometimes surrounded by a bracteal involucre. *Flowers* bisexual. *Sepals* 5, free, variable in size and shape, herbaceous or leathery, persistent in fruit. *Corolla* actinomorphic, infundibuliform or rarely hypocrateriform, purple, red, pink, white or yellow, rarely sky blue, throat often darker or lighter than the fauce. *Stamens* 5, unequal, two usually longer, or sometimes equal, inserted near the base of the tube, included or rarely exserted; pollen pantorporate, echinate; disc annular. *Ovary* 2 – 3-locular, ovules 4 (–6–10); style 1, filiform, generally glabrous, with 2 globose stigmas, less commonly slightly elongated. *Fruit* a capsule, ovoid or globose, dehiscing by 2 – 10 valves or by irregular tears, with (2 –) 4 (– 10) seeds. *Seeds* most commonly trigonous or globose, of various size, glabrous or of varying indumentum.

#### DISTRIBUTION

Genus comprises 635 species, widespread across all tropical regions; it is the most species-rich genus of Convolvulaceae in Tropical Africa (Mwanga-Mwanga, Sosef & Simões, 2022; POWO 2024). In Ghana, 28 species are present.

#### NOTE

The current conserved type of *Ipomoea* L. is *Ipomoea pes-tigridis* L., following a proposal by Manitz (1976). A later proposal to change the conserved type of *Ipomoea* L. to *Ipomoea triloba* L. has been submitted to the Nomenclature Committee (Eserman *et al.,* 2020; Eserman *et al.,* 2023); this proposal has been recommended by the Nomenclature Committee (Applequist, 2023) and the General Committee (Wilson, 2024), and will be submitted to voting to the Nomenclature Session at the International Botanical Congress 2024, after which, if approved, the change of conserved type of *Ipomoea* L. to *Ipomoea triloba* L. would be effective.

### 1. *Ipomoea intrapilosa* Rose (1894: 367)

> Type: Mexico, Jalisco, *E. Palmer* 703 (lectotype US! US00111405, isolectotypes BM, GH, K, MEXU).

Perennial *shrub* or *tree* 3 – 10 m high. *Stems* light coloured, up to 50 cm in diameter. *Leaves*: petiole pale green, 3 – 9 cm long, glabrous; leaf blade green, lanceolate to narrowly ovate, 7 – 14 x 3 – 5.5 cm, rounded, acuminate, entire, glabrous or sparsely pubescent on the lower surface near the base of the midrib, 10 – 17 lateral veins on each side of the midrib, sometimes bi-glandular at the base of the midrib. *Inflorescences* simple cyme, terminal or axillary, 1 – 3 (– 5) flowers, pedunculate, peduncle 0.4 – 2 cm long, glabrous; bracteoles ovate to oblong, 3 – 6 mm x 1 – 2.5 mm, glabrous on both surfaces and pubescent within, caducous. *Flower* pedicel 1.8 – 5 cm long, glabrous; calyx coriaceous, ovate to obtuse or obtuse-mucronate and acute at the apex, 13 – 19 mm x 7 – 13 mm, abaxial surface glabrous and adaxial surface with soft, straight appressed hairs; corolla white or yellowish white, the tube and the interplical regions greenish yellow, funnel, 5 – 8 cm x 5 – 7 cm, glabrous or sparsely pubescent along the margins of the inter apical regions; stamens included, filaments 3 – 4 cm long, anthers 8 – 10.5 mm long, basal hairs up to 2.5 mm long; ovary 2 – locular, 4–ovuled, style 3.5– – 4 cm long, stigmas 2, globose to slightly elongate, 1 mm long. *Fruit* capsule, opening by 4-valved, 2 – 2.5 cm long; *seeds* pilose, with long hairs on the margins, 10 – 15 mm long.

#### DISTRIBUTION

Native to Mexico, introduced, cultivated or escaped from cultivation elsewhere. In Ghana: Eastern region, Greater Accra region. **New record for Ghana.**

#### SPECIMENS EXAMINED. GHANA

Greater Accra region: Botany Department, University College, Achimota, 21 Nov. 1959, *G.K. Akpabla* 490 (GC); Eastern region: Aburi Botanical Garden, 25 Mar. 1976, *Hall, Lock & Abbiw* 45849 (GC).

#### HABITAT

Dry shrublands in central Mexico, oak forests and tropical deciduous woodlands; alt. 900 – 2200m.

#### CONSERVATION STATUS

LC: Least Concern (Salas *et al.,* 2020).

#### PHENOLOGY

Flowers and fruits between October and April.

#### VERNACULAR NAMES

“cazahuate, “cazahuate blanco” or palo blanco” (Spanish)

#### USES

Ornamental and medicinal. (Osuna *et al.,* 1996, Bah *et al*., 2007, Wood *et al.,* 2020).

### 2. *Ipomoea verbascoidea* Choisy (1838: 56)

> Type: Angola, *J.J. da Silva s.n.* (holotype P00150787!, isotype P00150788!).

*Shrub*, or stout liana, with an enlarged storage root. *Stem* sub – erect to climbing, subcylindrical, longitudinally striate, white or yellow tomentose in young state, becoming sparsely pubescent to glabrous with age. *Leaves* petiole 2 – 10 cm, pubescent, carrying 2 glands at the insertion of the leaf lamina; leaf largely ovate to suborbicular, 4 – 15 × 3 – 17 cm, apex rounded to retuse and shortly apiculate, base cordate to truncate, margin entire to slightly sinuate, adaxial face finely pubescent, abaxial face densely floccose tomentose, white to greyish. *Inflorescences* axillary lax cymes, 1 – 3 flowered; peduncle 1 – 3cm, tomentose as the stem; bracteoles slightly unequal, linear – oblong, up to 2cm long, tomentose, carinate.

*Flower* pedicel c. 1cm long, pubescent; sepals subequal, broadly elliptic, coriaceous, pubescent; outer ones 11 – 16 × 6 – 8 mm, inner ones 13 – 17 × (5 –)7 mm; corolla broadly tubular, 6 – 12cm long, pink, mauve or white, with darker purple center, glabrous; stamens of equal length, up to 6 cm long, widened and pubescent at the base, anther narrowly obolid, 6 – 12 mm long, sagittate at the base; ovary ovoid, 2.5 – 3mm long, 2-locular, 4-celled, glabrous; style filiform, 4.6 – 8 (– 8.5) cm long; stigmas 2, globose. *Fruit* capsule, opening by 4-valves, oblong ovoid, coriaceous, glabrous, completely enclosed by the persisting sepals, coriaceous; seeds brown, ovoid, 6 – 8mm densely pubescent, covered in long, cottony, white hairs.

#### DISTRIBUTION

Native to and occurs in Angola, Botswana, Cameroon, Caprivi Strip, Congo, Ghana, Guinea, Ivory Coast, Madagascar, Malawi, Mozambique, Namibia, Nigeria, Sudan, Tanzania, Togo, Uganda, Zambia, Zaïre, Zimbabwe. In Ghana: Northern region and Brong Ahafo region (Fig. 1).

**Figure 1.**
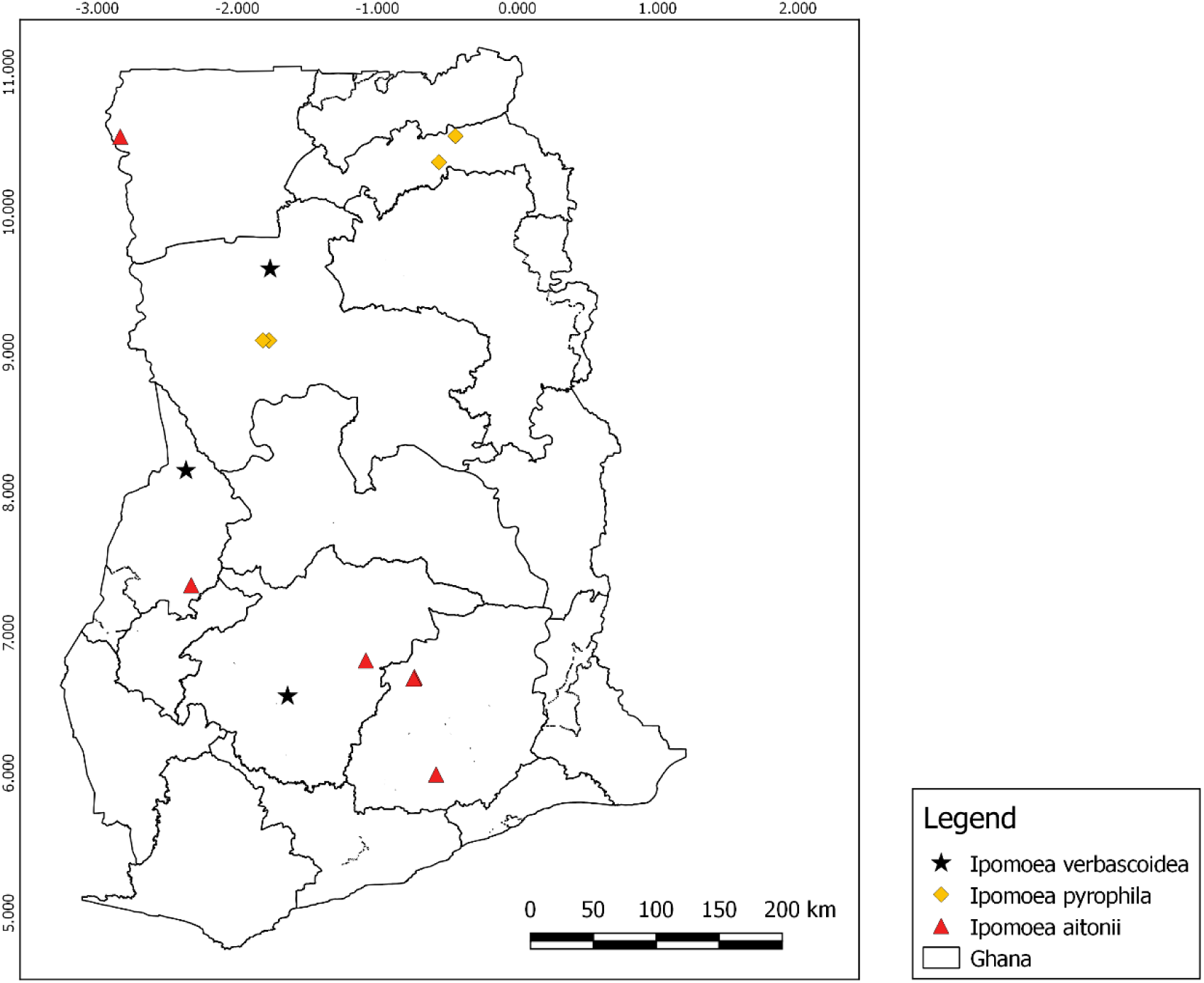
Distribution records of *I. verbascoidea*, *I. pyrophila* and *I. aitonii* in Ghana.

#### HABITAT

Often found growing in woodlands, open grassland, at elevations from 490 – 1,525m; some populations can be found in forests, also in subhumid, dry, subarid environments.

#### CONSERVATION STATUS (PRELIMINARY ASSESSMENT)

LC: Least Concern.

#### PHENOLOGY

Reported to flower at different times of the year in different countries; in Southern Africa, reported to flower from November to March; in Namibia, it flowers in midsummer (Prota4U.org).

#### VERNACULAR NAMES

*damara creeper* (English); *damarawinde* (German). In Namibia, known by several names: *’hoan* (Ju), *n!’huru* (Kung bushmen); *engamukuiju* (Kwanyama); *engamukuiyu* (Oshikwanyama); *engamukuiyu* (Oshiwambo); otjindwapa (Otjiherero) (Verdcourt 1963, Prota4U.org).

#### USES

The root can be used to obtain refreshing or lifesaving water (to quench thirst): the root is grated, the shavings put in one hand, squeezed and the liquid let to run down along the thumb right into the mouth; however, the liquid is said to negatively affect one’s ability to move and as a result one is forced to wait until the side effects wear off, for which there may be a slightly narcotic or paralysing effect; the Tswana people of Southern African countries pound the root in milk to make a potion for anorexia and poor weight gain. It is also used as food for bucks, especially antelope species (Prota4U.org).

#### SPECIMENS EXAMINED. GHANA

Northern region: Konkori, Mole N.P, 28 Jul. 1976*, Hall & Lack* 46265 (GC).

### 3. *Ipomoea cairica* (L.) Sweet (1826: 287)

> *Convolvulus cairicus* L. (1759: 922).
>
> Type: Icon, t. 70, Vesling in Alpino, De Plantis Aegypti (1640)

Perennial, climbing, *herb*, with enlarged storage roots. *Stems* twining or prostrate, completely glabrous or shortly pubescent at the nodes, up to 2cm long, sometimes shallowly muricate. *Leaves*: petiole 2 – 6 cm long, terete, glabrous; leaf palmately compound, orbicular, ovate or elliptic in outline, 5 – 7 lobes, narrowly elliptic, elliptic or ovate, apex acuminate, base attenuate, outer lobes often bifid, glabrous, 3 – 10cm, apex acute or obtuse, mucronulate or attenuate; with pseudostipules, 5 – 7 palmately compound as the leaves, glabrous, at the insertion of the petiole. *Inflorescences* in lax dichotomous cymes, 1 to many flowered; peduncle 5 – 8 mm long, ramified; bracteoles 1.5 mm long, caducous. *Flower* pedicel up to 3 cm long, thickening towards the apex; sepals subequal, ovate, 4 – 6.5 × 2.5 – 3 mm, glabrous, sometimes verruculose, the outer ones slightly shorter, obtuse to acute and mucronulate at the apex, membranous and pale along the margins; corolla funnel-shaped, (3 –) 4 – 6.5 × 4 – 5 cm, pinkish-mauve, purple, to entirely white, with a purple center, glabrous; stamens included, filaments unequal, dilated and pubescent at the base, anthers 2.5 mm long, sagittate at the base, pollen pantoporate, spinulose; ovary 2-locular, style included, c. 18 mm, glabrous. *Fruit* subglobose capsule, glabrous, 1 – 1.2 cm in diameter; *seeds* subglobose or ovoid, 4.2 – 6 mm long, blackish, densely short-tomentose with long silky hairs along the edges.

#### DISTRIBUTION

Native in Tropical and South Africa, West Indian Ocean, Arabian Peninsula to Temperate East Asia. In Ghana: Greater Accra region (Legon, Medea), Central (Elimina), Eastern (Suhum) and Northern regions (Bole) (Fig. 2).

**Figure 2.**
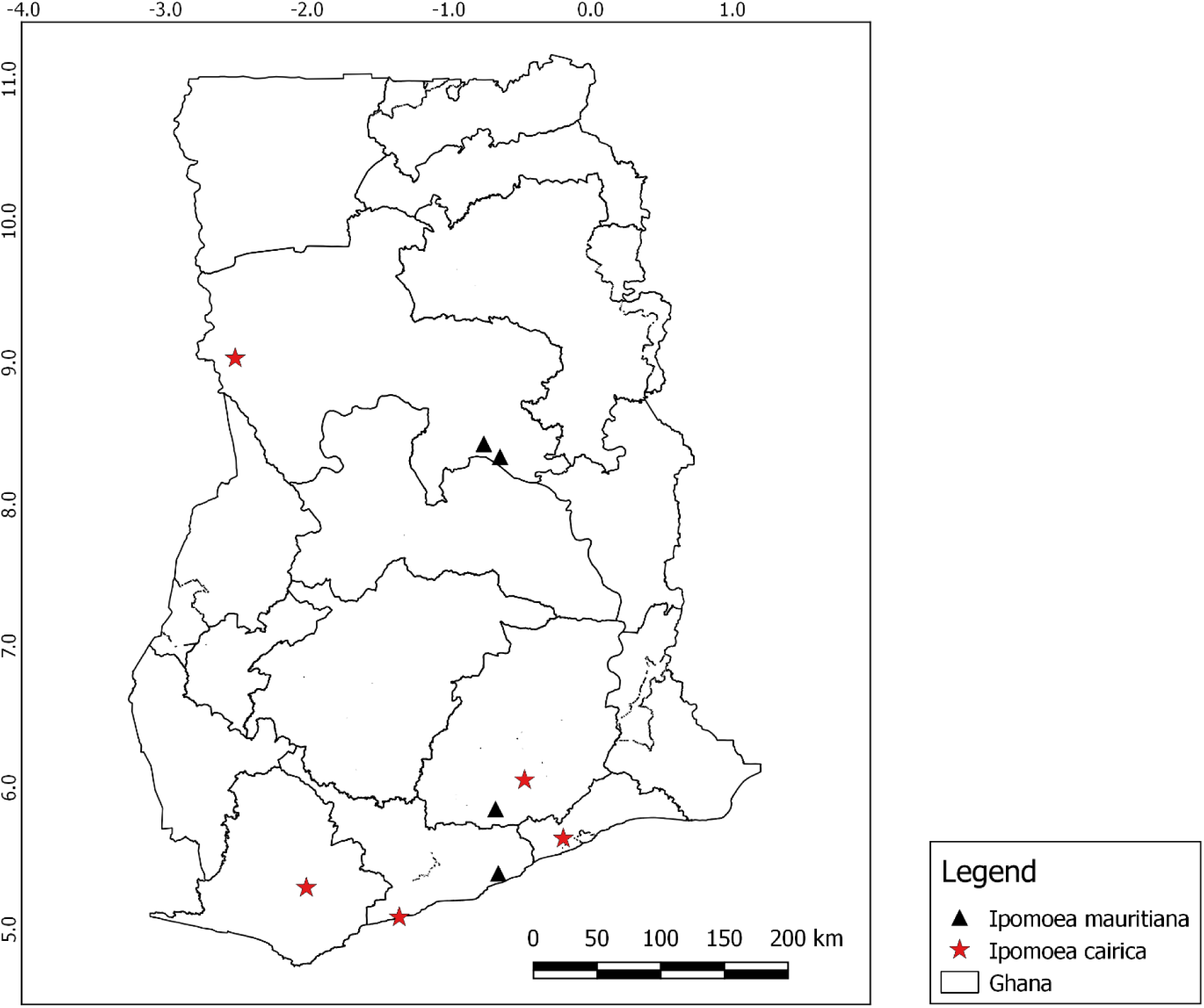
Distribution records of *I. mauritiana* and *I. cairica* in Ghana.

#### HABITAT

Grows primarily in the seasonally dry tropical biome; secondary forest and savannah (POWO, 2024).

#### CONSERVATION STATUS

LC: Least Concern (Allen, 2017).

#### USES

Most parts of *Ipomoea cairica* have been recorded to be edible; the leaves are eaten when still young, and roots are cooked before consuming. Leaves of the species are also used for treating skin disorders and seeds are used as laxatives, purgatives and enemas. It is also used medicinally by Zulu people, who make a mixture using crushed leaves and drink it for healing body rashes and fever (Burkill, 1985).

#### SPECIMENS EXAMINED. GHANA

Greater Accra region: Legon hill, 15 Jan. 1956, *C.D. Adams,* 3700 (GC); Central region: Elimina-Emisano, 1 Dec. 1929, *L.O*. *Deakun* 53 (GC); Eastern region: Suhum, 1 Dec. 1951, *J.K. Morton* 6333 (GC); Northern region: Bole Banbow, Bole, 1 Nov. 1958, *T.M. Hams* s.n. (GC).

### 4. *Ipomoea mauritiana* Jacq.(1791: 216)

> Type: Jacquin, Pl. Hort. Schoenbr.: tab. 200 (1797).

Annual, climbing, *herb,* with enlarged storage roots*. Stems* prostrate, often woody in the lower parts, cylindrical, more or less fistulose, muricate, with faint longitudinal ridges, ramified, glabrous. *Leaves*: petiole 3 – 15 cm long, glabrous or muricate; leaf deeply palmately lobed, sub – orbicular in outline, 7 – 18 cm x 6 – 17 cm, base cordate, apex acuminate or obtuse and mucronulate, 5 – 7 lobed, the outer lobes sometimes bifid, margin entire, overall glabrous except puberulent below, near the base or sparsely pubescent along the main veins. *Inflorescences* few to many-flowered cyme, puberulent; peduncle cylindrical or slightly angular, 2.5 – 20 cm long; bracteoles oblong, caducous, 1 – 2 mm. *Flower* pedicel cylindrical, 1.5 – 4.5 cm long, glabrous; sepals subequal, suborbicular to broadly elliptic, 6 – 7 mm long, imbricate and appressed to the corolla tube, rounded to obtuse at the apex, coriaceous, margin hyaline, glabrous; corolla funnel-shaped, 5 – 8 cm long, bright pink to whitish, with a purple darker center, glabrous, with a broad cylindrical tube, gradually opening up towards the apex, often with lobes deeply marked and spreading; stamens included, filaments up to 2 cm long, widened and pubescent at the base, anthers 3 – 4 mm long, white, sagittate at the base; ovary conical, 2-locular, glabrous; style filiform, puberulous. *Fruit* capsule, ovoid, 10 – 15 × 8 – 15 mm obtuse at the apex, 2-locular, glabrous; seeds 5 – 7 mm, black, covered in long white silky hairs.

#### DISTRIBUTION

Native in Tropical America and Africa, introduced in other tropical and subtropical regions (POWO, 2024). In Ghana, widespread: Greater Accra, Ashanti, Brong Ahafo, Volta, Eastern, Northern and Central regions (Fig. 2).

#### HABITAT

Growing primarily in the seasonally dry tropical biome; scattered throughout the tropics but rarely abundant (POWO, 2024).

#### CONSERVATION STATUS (PRELIMINARY ASSESSMENT)

LC: Least Concern.

#### VERNACULAR NAMES

*finger-leaf morning-glory* (English) (Prota4U.org).

#### USES

The plant is extensively used across Asia, especially for medicinal purposes: in Peninsular Malaysia, the root is pounded and applied to swellings; in India and the Philippines, the root is considered tonic, alterative, aphrodisiac, demulcent, lactagogue and cholagogue, and is used for fever and bronchitis; the powdered root is given for diseases of spleen and liver, for menorrhagia, debility and fat accumulation. In India, the seeds are also used for coagulating milk. The plant is also grown for ornamental purposes, and as a fodder for cattle (Prota4U.org).

#### SPECIMENS EXAMINED (SELECTED). GHANA

Greater Accra region: Legon Hill, 11 Dec. 1955, *C.D. Adams* 3587 (GC); Ashanti region: Atewa Range F.R., alt. 2600’, 13 May 1967, *Enti & Jenik* 36454 (GC); Brong Ahafo region: Yeji on the Tamale road, 13 Aug. 1963, *M. Ansa-Emmim & S.K. Adom-Boafo* 245 (GC); Volta region: Chai River Forest Reserve [8°06’N, 0°24’E], Alt 400 m, 22 May 1996, *C.C.H. Jongkind & C.M.J. Nieuwenhuis* 2806 (GC); Eastern region: Asamankese roadside, 1 Jan. 1929, *E.D. Plumptre* 72 (GC).

#### NOTES

An American plant now widely distributed in the tropics of the Old World; naturalized in waste places and residential areas.

### 5. *Ipomoea aquatica* Forssk. (1775: 444)

Type: Yemen, Zabid, 5 Apr. 1763, *Forsskål 447* (holotype C10002419!, isotype BM).

Perennial, less often annual, fleshy, stoloniferous, *herb*. *Stems* prostrate or floating, fistulose, rooting at the nodes, terete, glabrous or pilose at the nodes, smooth, with longitudinal ridges. *Leaves* petiole 2.5-25 cm long, glabrous; leaf simple, linear, lanceolate, ovate to broadly triangular in outline, 3 – 13 (– 17) cm x 0.5 – 9 cm, base cordate, sagittate or hastate, apex acute, acuminate, margin entire, glabrous, or more rarely pilose. *Inflorescences* few-flowered cyme; peduncle up to 14 cm long, thinner than the petiole, glabrous; bracteoles scale-like, narrowly ovate, 1.5 – 2 mm long, apex acute. *Flower* pedicel 1.5 – 6.5 cm long, glabrous or puberulent at the base; sepals subequal, the outer ones slightly shorter than the inner ones, pale along the margins, glabrous, ovate, 7 – 12 mm long, apex obtuse, often mucronulate, inner sepals ovate-elliptic, c. 8mm long, apex mucronate; corolla funnel – shaped with a narrow tube, (2) 4.5–10 cm long, pink, lilac, pale, dark red, purple with a purple center, or rarely white with dark purple center, glabrous; stamens unequal, longer ones up to 10 mm, shorter ones 4 – 5mm, widened and puberulent at the base, anthers 2mm long, sagittate; ovary obpyriform, 2 – locular, glabrous; style up to 13mm long, articulated. *Fruit* capsule, tardily dehiscent or indehiscent, ovoid to globose, 10 – 14 mm long, 8 – 12 mm wide; *seeds* ovoid, densely pubescent.

#### DISTRIBUTION

In Tropical and subtropical regions of Africa and Asia. In Ghana: **N**orthern, Brong Ahafo, Central, Upper West, Greater Accra and Volta regions.

#### HABITAT

A helophyte, growing primarily in the seasonally dry tropical biome; in moist, marshy or inundated localities, shallow pools, ditches, rice fields, forming dense masses; also found along roadsides, at elevations from sea-level up to 1,000 m.

#### CONSERVATION STATUS

LC: Least Concern (Gupta & Sayer, 2018).

#### VERNACULAR NAMES

water-spinach (English); *weng cai*, *kangkong* (China), *liseron d’eau, patate aquatique* (French), *marol* (Somali), *bhaji, karmi bhaji, marmi bhaji* (India), *swamp cabbage* (Trinidad & Tobago) (Dueñas-López, 2023).

#### USES

The leaf sap of *I. aquatica* is used in medicine for treating insanity; the young plants and leaf are eaten as food either cooked or uncooked; whole plant of *I. aquatica* is used in medicine for general body healing and the flower buds is used for treating skin diseases (Burkill, 1985; Van Wyk, 2005).

#### SPECIMENS EXAMINED (SELECTED). GHANA

Greater Accra Region: Mile 12 Accra-Winneba road [5° 34’25”N, 0°15’04”W], 10 Jun. 1961, *Hall* 1873 (GC); Central region: Cape Coast, marshy roadside, 15 Dec. 1964, *J.B. Hall* 2775 (GC); Northern Region: Mole Game Reserve [9°29’55”N, 1°59’55”W], 21 Nov. 1994, *C.C.H. Jongkind & C.M.J. Nieuwenhuis* 1891 (GC); ca. half mile North of Yeji [8°12’50”N, 0°38’42”W], 13 Aug. 1963, *Ansah-Emmim* & *Adom-Boafo* VBS 233 (GC); Yeji, [8°12’50”N, 0°38’42”W], 11 Apr. 1964, *Hall* VBS 1277 (GC).

### 6. *Ipomoea asarifolia* (Desr.) Roem & Schult. (1819: 251) *Convolvulus asarifolius* Desr. (1792: 562)

Type: Senegal, *Roussillon s.n.* (holotype P-LA[P-LAM00357544]; isotype P-JUSS[P-JUSS6798]).

Perennial *herb*. *Stems* prostrate, sometimes twinning, terete or angular, puberulent, with longitudinal ridges. *Leaves* petiole 3 – 8.5 (– 13) cm long, glabrous, thickened and with longitudinal ridges, or minutely muricate; leaf simple, circular to reniform, 3.5 – 7 (– 13) cm x 3.5 – 8.5 (– 18) cm, apex obtuse to emarginate, base cordate with rounded lobes, margin entire, glabrous, subcoriaceous. *Inflorescences* a lax cyme, 1 – few (– many) flowered; peduncle 2 – 5 (– 9.2) cm; bracteoles ovate, lanceolate, minute, 1 – 2 mm long; pedicel 1 – 3 cm long, glabrous; sepals unequal, elliptic – oblong, apex obtuse, mucrunolate; outer ones shorter, 5 – 8 mm long, more or less muricate, inner ones longer, 8 – 11mm long. *Flower* corolla funnel-shaped, reddish purple, 6 – 8cm long, glabrous. *Fruit*: capsule, globose, 8.5 –10.2 mm long; seeds not seen.

#### DISTRIBUTION

Widely distributed across tropical regions (POWO, 2024). In Ghana: Upper East, Eastern, Upper West, Greater Accra, Northern and Central regions (Fig. 3).

**Figure 3.**
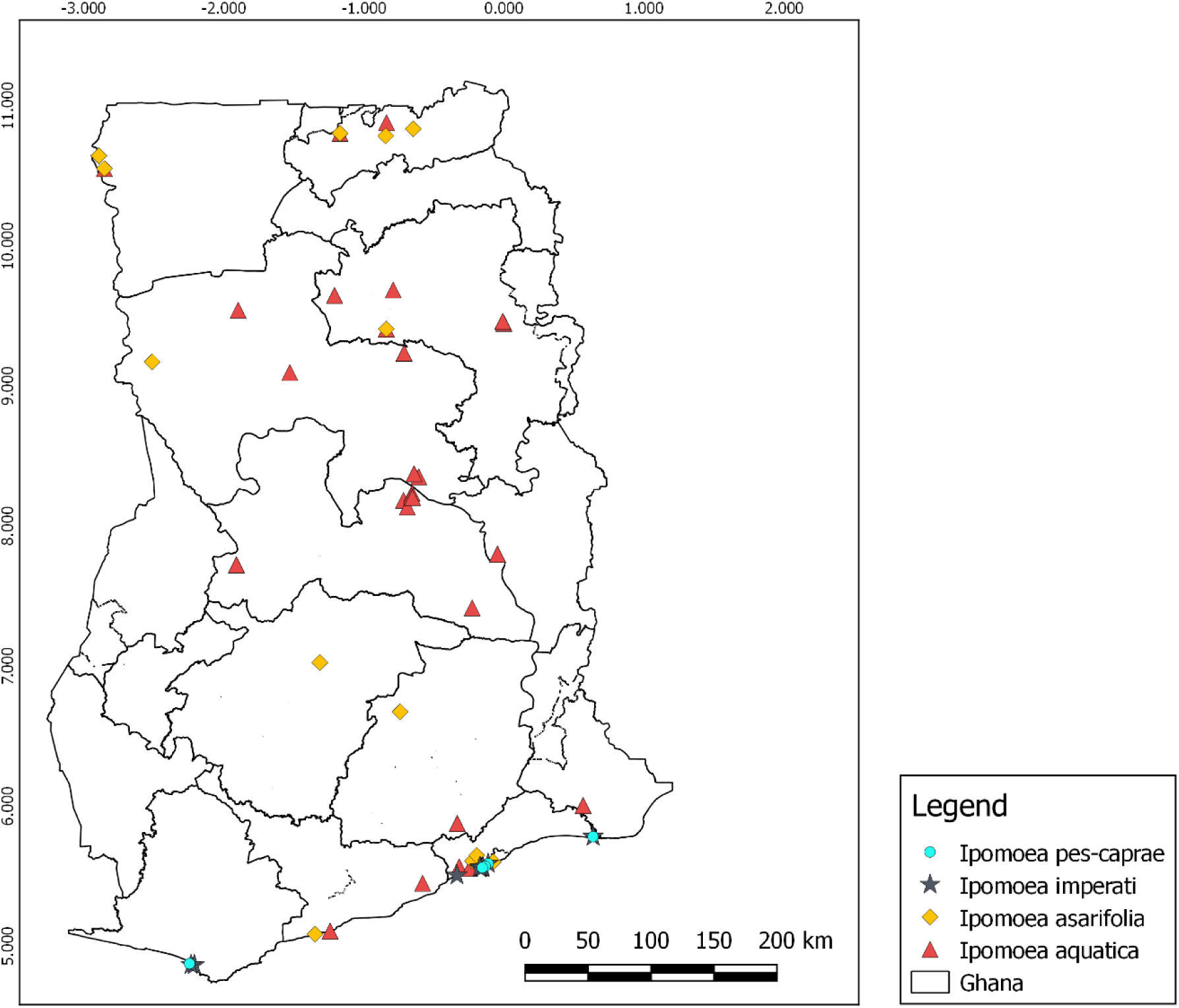
Distribution records of *I. pes-caprae*, *I. imperati*, *I. asarifolia* and *I. aquatica* in Ghana.

#### HABITAT

A scrambling herb, growing primarily in the seasonally dry tropical biome (POWO, 2024); in sandy areas and waste places in Africa; in marshy grasslands and waysides at elevations up to 250m (Prota4U.org).

#### CONSERVATION STATUS (PRELIMINARY ASSESSMENT)

LC: Least Concern.

#### USES

Whole plant is used for treating arthritis, rheumatism, lumbago, eye treatment; leaves are used for treating cutaneous and subcutaneous parasites (Burkill, 1985); the leaves are cooked and eaten as a vegetable, also used as dye, tying material and tinder, purgative, abortifacient (Prota4U.org).

#### SPECIMENS EXAMINED. GHANA

Eastern Region: Bumso [6°17’19”N, 0°28’39”W], *s.c.* s.n. (MO); Upper East Region: Bals of White Volta on road from Bolgatanga to Bawku [10°47’36”N, 0°51’38”W], 4 Apr. 1953, *Morton* s.n. (GC); Before Yamlapa, 22 May 1952, *Morton* s.n. (GC); Busufo grassland, 19 Dec. 1950, *Adams* & *Akpabla* s.n. (GC); Yamlapa, 1 Jun. 1958, *Harris* s.n. (GC); Northern region: Old Dam at Tamale, 18 Jan. 1966, *A.A.Enti & C.W. Agyakwa,* VBS 468 (GC).

### 7. *Ipomoea imperati* (Vahl) Griseb. (1866: 203)

> *Convolvulus imperati* Vahl (1790: 17).
>
> Type: Imperato, Hist. Nat., éd. 2: 671 (1672)

Perennial, fleshy, stoloniferous, *herb*. *Stems* prostrate, sometimes twining, terete, glabrous, rooting at the nodes. *Leaves* petiole 0.5 – 4.5 (– 5) cm, glabrous or less often puberulent at the apex; leaf simple, very variable in shape and size, particularly on the same plant, linear, lanceolate to narrowly elliptic-oblong or ovate, sometimes shallowly 3 – 5-lobed, 1.5 – 14 × 1.5 – 3.5 (– 6.5) cm, apex bilobed, emarginate, obtuse to acute or mucronate, base rounded to truncate or cordate, margin entire to undulate. *Inflorescences* lax cymes, 1 – 3-flowered; peduncle 0.6 – 1.9 cm; bracteoles very narrowly elliptic-oblong to linear or subulate, 2 – 4 mm long. *Flower* pedicel 2.1 – 4.3 cm; slightly club-shaped; sepals unequal, oblong, obtuse to shortly cuspidate at the apex, subcoriaceous, outer ones shorter, 8 – 9 mm long, inner ones longer, 10 – 11 mm long; corolla funnel – shaped, 3,9 – 5 cm long, white, yellowish on the inside, with a dark purple center, glabrous; stamens unequal, longer filaments 8 – 10 mm, shorter filaments 5 – 6 (– 7) mm, puberulent at the base, anthers pale yellow, 3 – 4 mm long; ovary ovoid, 2-locular, 2.2 – 2.5 mm high, glabrous; style filiform, 12 – 13 mm long. *Fruit* capsule, ellipsoid to sub-globose, dehiscing by 4 valves, glabrous; seeds ovoid-trigonous, 4 – –5mm long, tomentose with greyish silky hairs.

#### DISTRIBUTION

Tropical and subtropical coasts (POWO, 2024). In Ghana: Greater Accra and Western regions.

#### HABITAT

*I. imperati* grows on coastal flats, beaches, windward and leeward slopes of dunes, up to 100m (CABI, 2021).

#### CONSERVATION STATUS (PRELIMINARY ASSESSMENT)

LC: Least Concern.

#### USES

The plant is used in folk medicine for the treatment of inflammation, swelling and wounds, as well as to treat pains after childbirth and for stomach problems (De Paula–Zurrón *et al.,* 2010).

#### VERNACULAR NAMES

*beach morning-glory* (English) (CABI 2019).

#### PHENOLOGY

Flowering from May to August (GBIF, 2024).

#### SPECIMENS EXAMINED. GHANA

Greater Accra region: Accra, 2 May 1966, *T.W. Porown* 399 (GC); Medea, 21 May 1953, *D.W. Woodall* 15588 (GC); Labadi, 16 Oct. 1963, *J.K. Botokro* 47356 (GC); Accra Sea Coast, 13 Oct.1899, *T.W. Brown* 399 (GC); Near beach W of Accra, [5°30’06°N, 0°20’42’’W], alt 5m, 26 Nov. 1994, *C.C.H. Jongkind, D.K. Abbiw & C.M. Markwei* 1898 (GC).

### 8. *Ipomoea pes-caprae* (L.) R.Br. (1818: 477)

> *Convolvulus pes-caprae* L. (1753: 159).
>
> Type: Rheede, Hortus Ind. Malab. 11: t. 57.

Perennial *herb*, woody at the base, glabrous, containing an abundant white sap. *Stem* stoloniferous, prostrate, fistulose, often purplish, glabrous to slightly puberulent. *Leaves* petiole 2–17 cm long, purplish, glabrous; leaf erect, slightly succulent, suborbicular to square-shaped, 3 – 10.5 cm x 3 – 12 cm, apex emarginate and mucronate or truncate, sometimes lobed, base cuneate to rounded or cordate, glabrous or very sparsely puberulent on both sides, margins entire. *Inflorescences* lax cymes, terminal or axillary, 1 – 3 (– many) flowered; peduncle angular or flattened, 3 – 16 cm long, glabrous; bracteoles caducous, elliptic-ovate to narrowly elliptic – ovate, 3 – 3.5 mm. *Flower* pedicel 1.2 – 4.5 cm long, glabrous; sepals unequal, glabrous, the outer ones ovate to slightly elliptic, mucronate, 6 – 10mm long, distinctively 3 – 5 nerved, the inner ones larger, 8 – 15mm long; corolla funnel-shaped, pink or red purple with darker purple centre, 3 – 6 cm long, glabrous; stamens unequal, 5 – 9mm long, widened and pilose at the base; anthers 4 – 4,5 mm long, sagittate at the base; ovary globose to ovoid, 2 – 2.5 mm, 2-locular, glabrous, style 12 – 20 mm, stigmas 2, globose. *Fruit* capsule, globose, 12 – 18 mm long, 4-seeded or more, opening by 4 valves, glabrous; seeds black, trigonous-globose, 6 – 10 mm long, densely brownish tomentose.

#### DISTRIBUTION

Tropical and subtropical coasts (POWO, 2024). In Ghana: Greater Accra and Western regions.

#### HABITAT

*Ipomoea pes-caprae* is a pioneer coloniser of sand-dunes occurring throughout the seaboard of tropical regions (Prota4U.org).

#### VERNACULAR NAMES

*bababarakora* (Senegal: Madingue), *npiiti* (Nigeria: Yoruba), *batatilla* (Dominican Republic), *bay hops, bay winders* (Bahamas), *bejuco de playa* (Puerto Rico), *boniato de playa* (Cuba), *patate lan mer* (Dominican Republic), *patate marron* (Haiti), *Goat’s foot Convolvulus* (English) (Prota4U.org).

#### USES

The leaves are used to treat skin diseases (Burkill, 1985). The leaves are eaten as vegetables in Zanzibar, and also used for fodder/forage for pigs and cows in China; pulped leaves are rubbed on fishing nets in Malawi as a lure to entice the fishes to enter. Extract from the stems used for strong anti-tumour action; leaves used as anodynal in rheumatism and emollient on ulcerous and other sores, diuretic and laxative (Brown & Frank, 2023).

#### CONSERVATION STATUS

LC: Least Concern (Bárrios & Copeland, 2021).

#### SPECIMENS EXAMINED (SELECTED). GHANA

Greater Accra region: Near Labadi Lagoon, Shrub vegetation, 1 Jan. 1933, *F.R. Irvine* 1962 (GC); Labadi beach, 23 Jan. 1954, *W. Woodall* 16640 (GC); Western region: Beach near Axim, Shrub vegetation, 1 Mar. 1934, *F.R*. *Irvine* 2565 (GC).

### 9. *Ipomoea eriocarpa* R.Br. (1810: 484)

> Type: Australia, New Holland, *Banks & Solander s.n.* (holotype BM001040629!).

Annual *herb*, on a woody base. *Stems* prostrate or climbing, slender, up to 1 cm in diameter, pubescent to hispid, with a mixture of long and short hairs. *Leaves*: petiole 1 – 6 cm long, pubescent; leaf simple, ovate to widely cordate and narrowly linear to oblong-mucronate, 2 – 10 x 0.8 – 7 cm, base sub-hastate with rounded lobes, apex long attenuate to acuminate and mucronate, pilose to strigose, hairs highly concentrated on the midrib and the veins. *Inflorescences* axillary cymes, clusters of 3 – many flowers, rarely 1, subsessile or short pedunculate; peduncle 1 – 10mm long, densely pubescent; bracteoles linear to narrowly ovate-elliptic, 3 – 8mm, pubescent. *Flower* pedicel ca 5 mm long, pubescent; sepals subequal, ovate to lanceolate, apex acuminate, 6 – 8mm long, hispid to pilose, ciliate, spreading in fruit, inner sepals slightly narrower; corolla tubular to funnel-shaped, pink-purplish or rarely white, 6 – 9 x 13 mm, midpetaline areas pilose; stamens included, filaments inserted at the base of the corolla, glabrous, anther globose, 1mm long; disc 0,4 mm high, pubescent; ovary obpyriform, 2,5 mm high, hirsute, 2-locular, 4-celled; style distinctly articulate, 4 mm long; stigma 2-globose. *Fruit* capsule, ovoid-globose to globose, 5 – 6 mm in diameter, pubescent, apiculate, with persistent style; seeds black-grey, trigonous-globose, 2.5 mm long, glabrous, finely punctate.

#### DISTRIBUTION

Native to tropical regions in Africa, Australia, Asia and Saudi Arabia. Introduced in Oman, Puerto Rico, Réunion, Windward Is., Yemen (POWO, 2024). In Ghana: Upper West, Northern, Eastern, Brong Ahafo, Eastern and Volta regions (Fig. 4).

**Figure 4.**
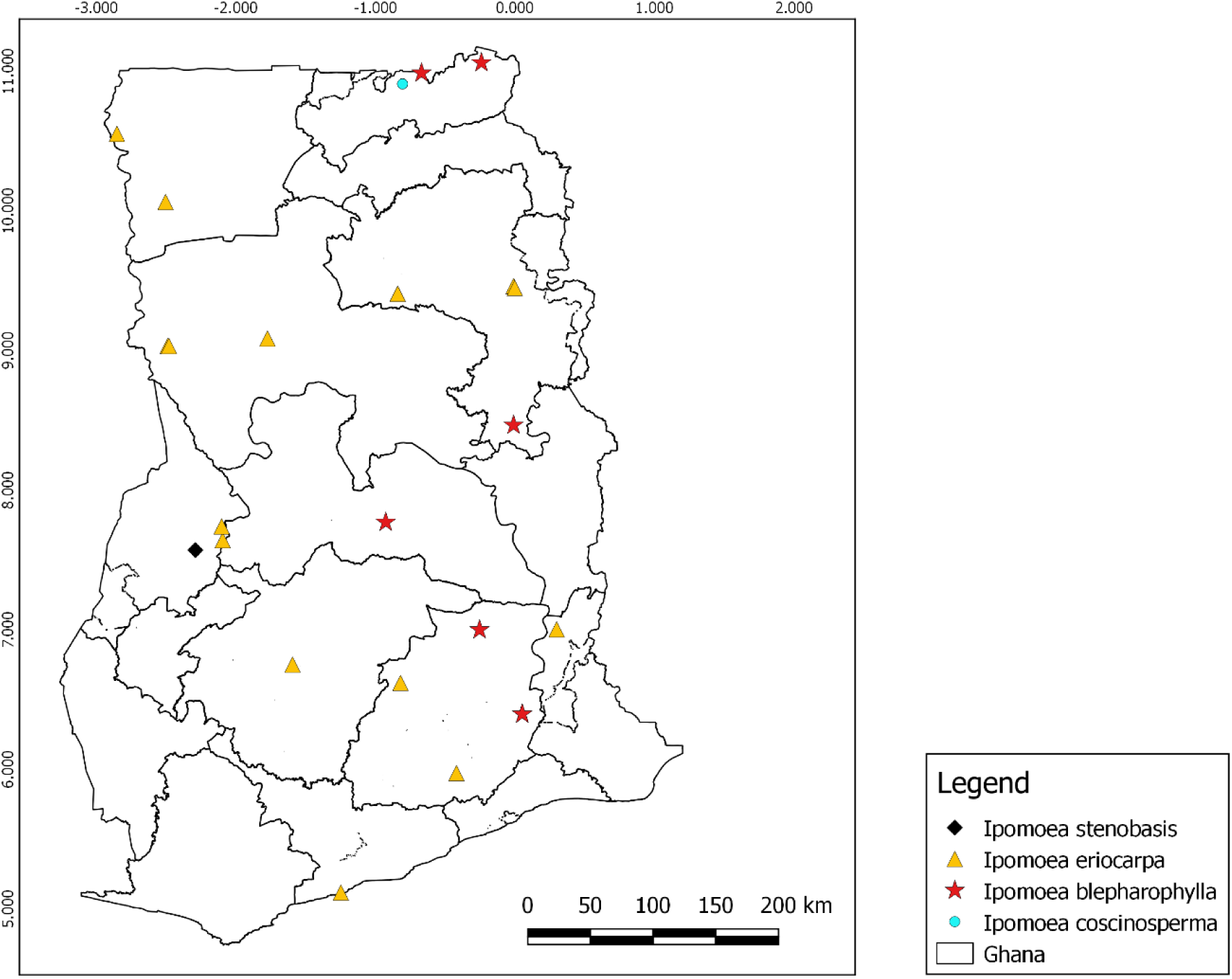
Distribution records of *I. stenobasis*, *I. eriocarpa*, *I. blepharophylla* and *I. coscinosperma* in Ghana.

#### HABITAT

Reported to be growing on grassland, hedgerows, waste spaces, cultivated ground, often on clay soils at altitudes of 0 – 1,350 m (Verdcourt, 1963); in Ethiopia, also in woodland habitat (*Combretum*-*Terminalia*-*Anogeissus* woodland), up to 1,700m (Demissew 2006).

#### CONSERVATION STATUS (PRELIMINARY ASSESSMENT)

LC: Least Concern.

#### VERNACULAR NAMES

*manding-bambara gabi* (Senegal), *manding-mandinka* (Gambia), *mende kpokpo* (Sierra Leone), *tiny morning glory* (English) (Prota4U.org).

#### USES

Seed oil from *Ipomoea eriocarpa* is used to treat skin diseases and dermal eruptions, arthritis and rheumatism (Burkil, 1985). Used in soups or mixed with other foods including other *Ipomoea* species (Nigeria); in Tanzania and India, the leaves are boiled and eaten as vegetables; the seeds have unspecified medicinal use in Gambia; subsequently, the oil extract of the plant is used in India for external application in headache, rheumatism, leprosy, epilepsy, ulcers and fevers (Prota4U.org).

#### SPECIMENS EXAMINED (SELECTED). GHANA

Eastern region: Afram Mankrong F.R, 19 Dec. 1957, *C.D. Adams* 4951(GC); Mt. Ejuanema, Kwahu, alt. 2200’, 25 Dec. 1957, *C.D. Adams* 5145 (GC); Northern region: Pong Tamale, Veterinary Dept., 27 Nov. 1935, *G.K. Akpabla* 378 (GC); Yendi, 12 Dec. 1951, *Adams & Akpabla* 4053 (GC); Bole, 1 Nov. 1958, *T.M.Harris* 955.

### 10. *Ipomoea coscinosperma* Hochst. ex Choisy, (1845: 354)

> Type: Sudan, Kordofan, *Kotschy 17* (lectotype G, isolectotype WAG).

Perennial, or sometimes annual, *herb*. *Stems* several, stout, suberect to prostrate, up to 3 m long, glabrescent or pilose. *Leaves*: petiole 0.5 – 1.2 cm long; leaf simple, linear-lanceolate to oblong, 2.4 – 8 cm x 0.5 – 2 cm, base cuneate or obtuse, apex subacute to obtuse and mucronate, margin entire, glabrescent or pilose. *Inflorescences* cymose, 1 – to few-flowered sessile clusters; peduncles inconspicuous, up to 5 mm long; bracteoles pilose, linear, inconspicuous, c. 4 mm long. *Flower* pedicels inconspicuous, up to 5mm long; sepals subequal, ovate-lanceolate to lanceolate, 6 mm long, up to 1.2 cm in fruit, apex long-attenuate, covered with long, white hairs, with hyaline lower margins; corolla narrowly funnel – shaped, small, only slightly longer than the calyx, red or white, 5 – 8 mm long. *Fruit*: capsule globose, glabrous, apiculate, with style base persistent, 5 – 7.5 mm in diameter; seeds brown, shortly pubescent, 3 mm long.

#### DISTRIBUTION

Native to Botswana, Chad, Eritrea, Ethiopia, Ghana, Guinea, Kenya, KwaZulu-Natal, Mali, Mauritania, Namibia, Niger, Nigeria, Northern Provinces, Senegal, Somalia, Sudan, Swaziland, Tanzania, Uganda, Zambia, Zimbabwe (POWO, 2024). In Ghana: Upper East region (Fig. 4).

#### HABITAT

Grows primarily in the Seasonally Dry Tropical biomes (POWO, 2024).

#### PRELIMINARY CONSERVATION ASSESSMENT

LC: Least Concern (proposed here).

#### VERNACULAR NAMES

*Ñiñéni*, *manding-bambara* (Senegal) (Burkill, 1985).

#### USES

The whole plant parts are used in traditional Medicine and fodder for livestock (Prota4U.org).

#### PHENOLOGY

Flowering from December to May (Roux, 2003).

#### SPECIMENS EXAMINED. GHANA

Upper East Region: Near Bongo, on route to Nangodi, N.T.S, 15 Nov. 1959, *J.K. Morton*, A 3804 (GC).

### 11. *Ipomoea blepharophylla* Hallier *f*. (1893: 125)

> Type: Sudan, Gr Periba Ghattas, *Schweinfurth 1818* (P00434136!).

Perennial *herb*, on a woody rootstock. *Stems* several, prostrate, slender, densely hirsute with yellowish appressed hairs. *Leaves* petiole up to 1.5 cm long, pubescent; leaf lanceolate or narrowly oblong-linear, 0 – 8 cm x 1.3 – 2.5 cm, base rounded or subcordate, apex obtuse or mucronate, margin entire, nearly glabrous or with odd hairs on midrib above and margins and veins beneath, ciliate at the apex. *Inflorescences* axillary, flowers solitary; peduncle 1.5 – 2 cm long, pubescent; bracteoles narrowly ovate to lanceolate, unequal in length, ca. 3.5 mm long, pubescent. *Flower* pedicel up to 1 cm long, pubescent, longer in fruit; sepals unequal, apex acute, covered in long appressed hairs, outer ones shorter, up to 18 mm long and 4 mm wide, pubescent and ciliate, inner ones more narrowly ovate, slightly longer than the inner; corolla narrowly funnel-shaped, ca. 2,5 cm, thrice longer than sepals, pale or red-purple with darker lines and throat, distinctly narrower below the tube, with long white hairs at the apex of the midpetaline bands; stamens included, filaments slightly unequal, widened and pubescent at the base, anther ovoid, 2mm long, sagittate at the base; disc cup-shaped; ovary ovoid, glabrous, style included, stigmas 2-globose. *Fruit* capsule, globose, 9 – 10 mm long, glabrous, apiculate, with persistent style; seeds brownish, 4 – 4,5mm with appressed yellowish hairs, tomentose.

#### DISTRIBUTION

Angola, Benin, Burkina Faso, Burundi, Cameroon, Chad, Congo [Brazzaville], Ethiopia, Gabon, Ghana, Guinea, Ivory Coast, Kenya, Mali, Malawi, Mozambique, Nigeria, Rwanda, Senegal, South Sudan, Sudan, Tanzania, Togo, Uganda, Zambia, Zaïre, Zimbabwe (COL Checklist, 2022; POWO, 2024). In Ghana: Brong Ahafo, Upper East, Eastern and Volta regions (Fig. 4).

#### HABITAT

Grasslands (often seasonally flooded), wooded grassland after burning, secondary forests (dry evergreen) or rocky hills; (550-)1,080 – 1,860 m (Demissew, 2006; POWO, 2024).

#### CONSERVATION STATUS (PRELIMINARY ASSESSMENT)

LC: Least Concern.

#### SPECIMENS EXAMINED. GHANA

Upper East Region: Red volta F.R., Savannah, *Hall & Swaine* 46125, 24 Nov.1976 (GC); Upper East Region: Zowse, hills near Bawku, Grassland, *Hall & Enti* 35996, 8 Nov. 1966 (GC); Volta Region: 7 – 8m on Kete Krachi-atebubu road, *C.D. Adam* 4601, 19 Dec. 1956 (GC); Eastern Region: Jaketi on Afram plains, *G.K. Akpabla* 1876, 1 Mar. 1958 (GC); Upper West: Tumu resthouse, *J.K. Morton* 7564, 25 May 1952 (GC); Northern region: Kpandai Leprosarium, *J.B. Hall* 38753, 1 Aug. 1968 (GC).

### 12. *Ipomoea aitonii* Lindl. (1835: t. 1794)

> Type: Illustration type in Edwards’s Bot. Reg. 21: t. 1794 (1835).

Perennial *herb*. *Stems* prostrate or twining, strong-stemmed, densely covered with white and yellow spreading hairs. *Leaves* petiole hairy, 4 – 7 cm; leaf simple, 3 – lobed, rarely entire, orbicular in outline, bristly pubescent above, white hairy below, apex acute or acuminate, base cordate, 4.5 – 13 x 4.5 – 13 cm wide. *Inflorescences* lax or dense cymes 1 – many-flowered, densely clustered; peduncle 1.5 – 7 (– 15) cm long; bracteoles 7 x 2mm. *Flower* sepals lanceolate, sticky glandular, with spreading hairs, 12 – 22 mm long, 25 – 30 mm wide; corolla 1.2 – 1.7 (– 2) cm long, pink or mauve, with dark purple centre, pubescent on the upper portion of the midpetaline bands. *Fruit* capsule ovoid, sparsely pubescent, 8 mm high; seeds ovoid, pubescent (rarely quite glabrous), 4 – 5mm long, black, white tomentose.

#### DISTRIBUTION

The native range of *Ipomoea aitonii* is Africa, South Arabian Peninsula and Indian subcontinent. In Ghana: Brong Ahafo, Eastern, Ashanti region and Upper West regions (Fig. 1).

#### HABITAT

Riverine forest, thickets, clearing in bushland, becoming a weed of cultivated ground.

#### USES

The seed is used in medicine as laxative; the leaf is used as fodder in agri-horticulture (Burkil, 1985).

#### PRELIMINARY CONSERVATION ASSESSMENT

LC: Least Concern (proposed here).

#### SPECIMENS EXAMINED. GHANA

Eastern region: Abetifi, 20 Dec. 1939, *Scholes* 110 (GC); Kyibi, Apapam, 16 Dec.1953, *J.K. Morton* 8155 (GC); Ashanti region: Ashanti Akyim, Agogo, 28 Dec. 1927, *F.R. Irvine* 583 (GC); Brong Ahafo region: Sunyani, 1000ft, 18 Dec. 1954, *C.D. Adams* 2754 (GC); Northern region: Talense south, near Burufo, 20 Dec. 1950, *C.D. Adams* 4451 (GC).

### 13. *Ipomoea involucrata* P. Beauv. (1816: 52)

> Type: Nigeria/Bénin, Oware, *Palisot de Beauvois s.n.* (holotype G00023040!).

Annual or perennial *herb*. *Stems* creeping or climbing, densely hirsute with yellowish hairs, to glabrescent. *Leaves* petiole 1.3 – 8 cm long, pubescent; leaf simple, ovate, 2 – 8 × 1.5 – 7 cm, base cordate, apex attenuate – acuminate to obtuse, mucronate, base cordate, glabrous to densely tomentose on both surfaces. *Inflorescences* in dense heads, few – many flowered, enclosed within a bracteal involucre; peduncle 2 – 12 (– 16) cm, pubescent; outer bracteoles connate into large pubescent foliaceous boat-shaped involucre, with 2 cups, 3 – 6 cm long, 0.8 – 1.5 cm wide, green, pubescent, inner bracteoles small, linear – oblong to obovate, 1.5 – 2 × 0.2 – 0.4 cm, acute to aristate at the apex. *Flower* pedicel 1.5 – 3 mm, pubescent; sepals narrowly elliptic, glabrescent to densely pubescent, outer ones lanceolate and acute, 6 – 15 × 4 mm, inner ones ovate, shorter; corolla funnel –shaped, purple, rose – red, mauve, white or white with dark purple centre, 3 – 5 cm long, midpetaline bands pilose on the outside; stamens unequal, the longer ones 10 – 15 m, the shorter ones 5 – 7 mm, widened and pubescent at the base; anthers white, 1.7 – 2 mm, sagittate at the base; ovary ovoid, c. 1 mm, glabrous; style 8 – 13 mm, stigmas 2, white, globose. *Fruit* capsule, globose, opening by 4 valves, 6 mm wide, glabrous; seeds pubescent or glabrous, 3.5 – 4 mm long, blackish, glabrous to shortly pubescent.

**DISTRIBUTION**

The native range of *Ipomoea involucrata* is Tropical and South Africa (POWO, 2024). In Ghana: Eastern, Greater Accra, Ashanti, Western, Oti, Central and Volta regions (Fig. 5).

**Figure 5.**
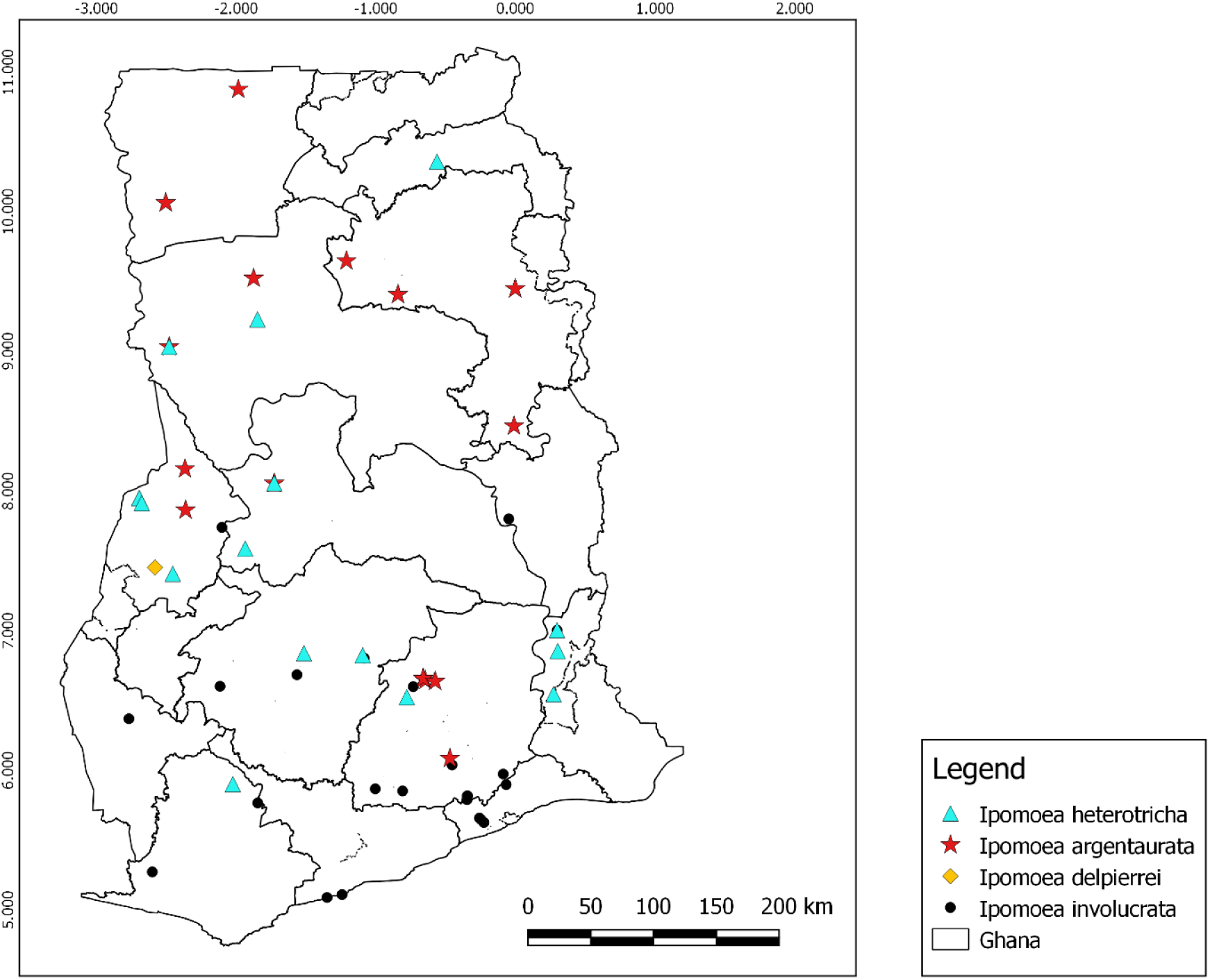
Distribution records of *I. heterotricha*, *I. argentaurata*, *I. delpierrei* and *I. involucrata* in Ghana.

**HABITAT**

Creepers on the roadside of the main road.

**CONSERVATION STATUS (PRELIMINARY ASSESSMENT)**

LC: Least Concern.

**USES**

The leaf sap is used as abortifacients and for the menstrual cycle; the seeds are used as laxatives, purgatives and enemas; the leaves are used in Medicine for treating pulmonary ailments; the leaves and roots are used as food, either cooked or uncooked (Burkill, 1985).

**SPECIMENS EXAMINED (SELECTED). GHANA**

Ashanti region: Atewa range [6°14’30’’N, 0°32’30’’W], 8 Jun. 1994, *C.C.H. Jongkind, D.K.Abbiw & C.M. Markwei* 1561 (GC); Western region: Amenfi east, Akropong, 25 Aug. 1898, *W.H. Johnson* 154 (GC); Ankasa Forest Resource Reserve, Ankasa based camp to Nkwanta guard camp, ca. 5.0 km. [5°13’06’’N, 2°39’06’’W], alt 150-230m, 13 Feb. 1999, *H.H. Schimdt, J. Stone, J. Amponsah & M. Chintoh* 3383(GC); Oti region: Kete Krachi, Mpuseto, 1 Dec. 1951, *J.K. Morton* 6430 (GC); Central region: Cape Coast, 30 May 1962, *J.B. Hall* 2328(GC).

### 14. *Ipomoea argentaurata* Hallier f. (1893: 132)

> Type: Central African Republic, Territoire du Chari, Krébedjé, Fort Sibut, Vallée de la moyenne Tomi, *Chevalier 5667* (syntypes P00434131, P00434129, P00434130).

Perennial *herb,* or sub-shrub. *Stems* prostrate or ascending from a woody base, densely whitish-tomentose with hispid yellow hairs. *Leaves* petiole 0.5 – 1.6 cm; leaf simple, oblong, ovate to lanceolate, 2 – 7 x 0.5 – 2.5 cm, base rounded, subcordate or cordate, apex attenuate, mucronate; densely strigose on the upper surface and deep green, silvery-silky below.

*Inflorescences* cymose, bracteate, axillary, multiple flowered; heads of flowers large, densely strigose, with long golden-yellow hairs; bracteoles 10 – 28 x 4 – 14 mm long, hairy like the calyx. *Flower* sepals linear-lanceolate or almost linear, acuminate, silky white on the back, with yellow strigose margin; corolla large, funnel-shaped, whitish turning light purplish with darker centre, c. 3 cm long, midpetaline bands strigose outside. *Fruit* capsule, 4-valved, glabrous; seeds covered with a dense dark brown pubescence.

#### DISTRIBUTION

Cameroon, Central African Republic, Chad, Ghana, Guinea, Ivory Coast, Nigeria, Sierra Leone, Togo, Benin, Guinea-Bissau, Burkina Faso, Mali and Senegal (Hassler 2022, POWO, 2024). In Ghana: Northern, Brong Ahafo, Eastern, Upper East and Upper West regions (Fig. 5).

#### HABITAT

A climber, growing primarily in the seasonally dry tropical biome; in savanna habitat at an altitude of up to about 300 m (Heine, 1963).

#### VERNACULAR NAMES

*ukpali*, *fárín gámó*, (Dagani, Dyokogye, Hausa, Ghana); *good luck* (English) (Burkill, 1985).

#### USES

The whole plant is used as genital stimulant or depressants; a decoction of aerial parts is drunk while kola nuts are eaten, in Ivory Coast, in the belief that it promotes spermatogenesis; other uses are mainly superstitious, as medicine for witchcraft, worn as an amulet, for example; clothing is fumigated with it, not as a scent, but as a charm for the same purpose, or for luck; in Bénin, a leaf decoction together with leaves of *Ficus vallischoudae* Delile, is drunk to treat hyperthermia and a decoction of the leafy twigs is taken to treat kwashiorkor (Burkill, 1985).

#### CONSERVATION STATUS (PRELIMINARY ASSESSMENT)

LC: Least Concern (proposed here).

#### SPECIMENS EXAMINED. GHANA

Northern Region: Damango, a yam farm near the Mole game reserve, Savannah grasslands, *A. A.* Enti, 6 Jan. 1965, 35162 (GC); Kwahu-Tafo, Grassland on flat rocks at 1500’, 19 Aug. 1963, *J.B. Hall* 0098 (GC); North, Yendi, 28 Dec. 1950, *C.D. Adams & G.K. Akpabla* 4051 (GC); Tamale, Tamale Girls School compound, *E*. *G. Asare*, 1 Oct. 1954, 5934 (GC); Kwahu-Abowom, on rocks by a pool, 12 Jun. 1970, *Hall & Agyakwa* GC 39684 (GC).

### 15. *Ipomoea heterotricha* F. Didr. (1854: 220)

> Type: Guinea, *Mortensen s.n.* (lectotype C10004084, isolectotype C10004083).

Annual *herb*. Stems twining, terete, densely covered in hirsute, long, yellowish indumentum, associated with a layer of greyish, shorter, hairs. *Leaves* petiole 1.7 – 3.4 cm long, densely hirsute as the stem; leaves simple, ovate to subtriangular or cordate, 4 – 7.8 x 2.4 – 6.1cm, base cordate to truncate, apex acute to acuminate, margin entire, densely yellowish pubescent above, with long silver dense hairs beneath. *Inflorescences* capituliform, dense, enlarged by persistent, enlarged, bracteoles; peduncle 2.9 – 4.6 cm, yellowish hirsute as the stem; bracts several, oval-elliptic to narrowly oblong, 15 – 30 × 5 – 8 mm, pilose as the leaves, the external face long hirsute. *Flower* sepals subequal, c. 9 mm long, shorter than the bracteoles, linear-lanceolate, apex acute, densely pubescent on both faces, strongly yellowish hirsute on the outside, as the bracts, towards the base, hirsute with silver hairs towards the apex; corolla funnel – shaped, white to pink with dark purple centre; stamens inserted, unequal, filaments 5 – 6 mm long, anther basifixed, ovoid, 1,8 mm long, sagittate at the base; ovary conical to ovoid, 2-locular, 5 – 6 mm long, glabrous; style filiform, 9 –13 mm. *Fruit* capsule, ovoid, 5 – 6 mm, glabrous; seeds ovoid, 3 – 4 mm, black, finely yellow pubescent.

#### DISTRIBUTION

Angola, Burundi, West Africa except Liberia, Central African Rep., Chad, Congo, Ethiopia, Kenya, Malawi, Rwanda, Senegal, Sudan, Tanzania, Uganda, Zambia, Zaïre, and Zimbabwe (POWO, 2024). In Ghana: Northern, Volta and Brong Ahafo regions (Fig. 5).

#### HABITAT

Found in open or dense *Combretum–Anogeissus leiocarpus* woodland or deciduous woodland; in Ethiopia, it can be found at 550 – 600 m altitude but extended to 435 – 1200 m altitude range in other countries; it can also be found in the undergrowth, woodland edge, wet grassland, rivers, and near waterfalls (Gonçalves, 1987).

#### PRELIMINARY CONSERVATION ASSESSMENT

LC: Least Concern (proposed here).

#### PHENOLOGY

Flowering from December to February, based on examined specimens.

#### ETYMOLOGY

The specific epithet “*heterotricha*” is gotten from the term “heterotrichy” which means different hair types (Wamucii, 2024).

#### SPECIMENS EXAMINED. GHANA

Brong-Ahafo Region: between Wenchi and Sunyani, 6 Feb.1995, *C.C.H. Jongkind* 2028 (GC); Brong-Ahafo Region: 5 Dec. 1995, *H.H*. *Schmidt, J*. *Amposah & A. Welsing* 1919 (GC); Eastern region: Mankrong, Kwahu, 23 Dec. 1957, *C.D*. *Adams* 5066 (GC); Northern region: Larabang-Wa road, ca 2 km [9°13’19’’N, 01°52’17’’W], alt. 140m, 1 Dec. 1995, *H.H*. *Schmidt, J*. *Amposah & A. Welsing* 1888 (GC).

### 16. *Ipomoea delpierrei* De Wild. (1911: 261)

> Type: Democratic Republic of Congo, Gumban (Uele), 1904, *Delpierre s.n.* (holotype BR0000008884404!).

*Herb*. *Stems* twining, striate, hirsute with long yellowish hairs. *Leaves* petiole 3 – 7.4 cm, densely long pubescent, grooved beneath, hirsute as the stems; leaf simple, subtriangular, ovate or cordate, 7.9 – 9.6 cm x 6.6 – 7 cm, base cordate, apex acute to acuminate, lateral nerves 13 – 14, margins entire to sinuate, densely long pubescent above and strigose beneath. *Inflorescences* capituliform, dense head of 3 – – 7 flowers; peduncle 11.8 – 13.5 cm, densely yellowish hirsute as the stem, enclosed by the bracteoles; bracteoles 2, foliaceous, ovate, 2 – 4.5 cm long, cordate at the base. *Flower* sessile; sepals unequal, shorter than the bracteoles, glabrous outside, denticulate and long ciliate inside, strigose, the outer sepals elliptic, 14 – 15mm long, inner ones narrowly oblong-elliptic, 12 – 13 mm long; corolla funnel-shaped, 3.5 – 5 cm, white with dark purple throat; midpetaline bands strigose on the upper portion; stamens pubescent throughout the length of the filaments, widened at the base, very unequal in length, longer ones 10 – 13.5 mm, shorter ones 2 – 4 mm, anther 3 – 4 mm, sagittate at the base, rounded at the apex; ovary ovoid, 1.5 mm high, glabrous, style filiform, 15 – 19 mm. *Fruit* capsule, globose, 7 mm in diameter, glabrous, 3 – 4-seeded; seed trigonous, ovoid or globose, 3 – 4 mm, dark brown, tomentose.

#### DISTRIBUTION

Native to Cameroon, Central Africa Republic, Ivory Coast, and Zaire (POWO, 2024). In Ghana: Brong Ahafo region (Fig. 5). **New record for Ghana**.

#### HABITAT

The species grows in woodlands and thickets (Verdcourt, 1963).

#### CONSERVATION STATUS (PRELIMINARY ASSESSMENT)

LC: Least Concern.

#### PHENOLOGY

Flowers in December.

#### ETYMOLOGY

The specific epithet “*delpierrei*” is named after the first collector, Delpierre (Wamucii, 2024).

#### SPECIMENS EXAMINED. GHANA

Brong Ahafo region: Injina road, Berekum, 12 Dec. 1954, *J.K*. *Morton* A1035 (GC).

### 17. *Ipomoea alba* L. (1753: 161)

> Type: Icon. *Munda-valli*, Rheede Hort. Ind. Malabar. 11: 103, t. 50 (1692).

Annual *herb*. *Stems* prostrate or twining, laticiferous, cylindrical, smooth, striate or sometimes muriculate, up to 5m long, glabrous or rarely pubescent. *Leaves* petiole 4.5 – 26 cm long, glabrous; leaf simple, entire to 3-lobed, ovate to orbicular in outline, or rarely ovate – oblong, 3 – 26 x 5 – 16 cm, apex acute to acuminate or obtuse, mucronate, base cordate, basal auricles rounded to angular, margin entire, covered in small blackish glands on both surfaces of the leaf. *Inflorescences* axillary, 1 – several-flowered; peduncle stout, 1 – 24 cm long; bracteoles small, caducous. *Flower* pedicels up to 3 cm long, thickening in fruit; sepals unequal, subcoriaceous, outer ones elliptic, 0.5 – 1.2 cm long with a long awn-like appendage, 6 – 9 mm long, often reflexed, inner ones 0.8 – 1.5 cm long, shortly acuminate in a triangular apex, 2.3 mm long, mucronulate; corolla hypocrateriform, cream-white, fragrant, night – flowering, tube 7 – 15 cm long, cylindrical to slightly angular, limb 11 – 16 mm wide; stamens slightly exserted, filaments subequal, 1 – 3 cm long, not widened at the base, inserted on the upper portion of the corolla tube, glabrous, anthers ovoid to obolid, 4 – 5mm long; ovary obpyriform, glabrous; style 10.5 – 12.2 cm long, glabrous. *Fruit* capsule, dehiscing by 4 valves, 2.5 – 3 × 1.5 – 2.3 cm, ovoid, mucronate, glabrous, 2-locular, apiculate; seeds 4, ovoid, 10 – 13 × 4 – 9 mm, brown or black, sparsely pubescent with white hairs.

#### DISTRIBUTION

Native to the Americas and introduced in Africa, Asia and Australia (POWO, 2024). In Ghana: Volta and Greater Accra region (Fig. 6).

**Figure 6.**
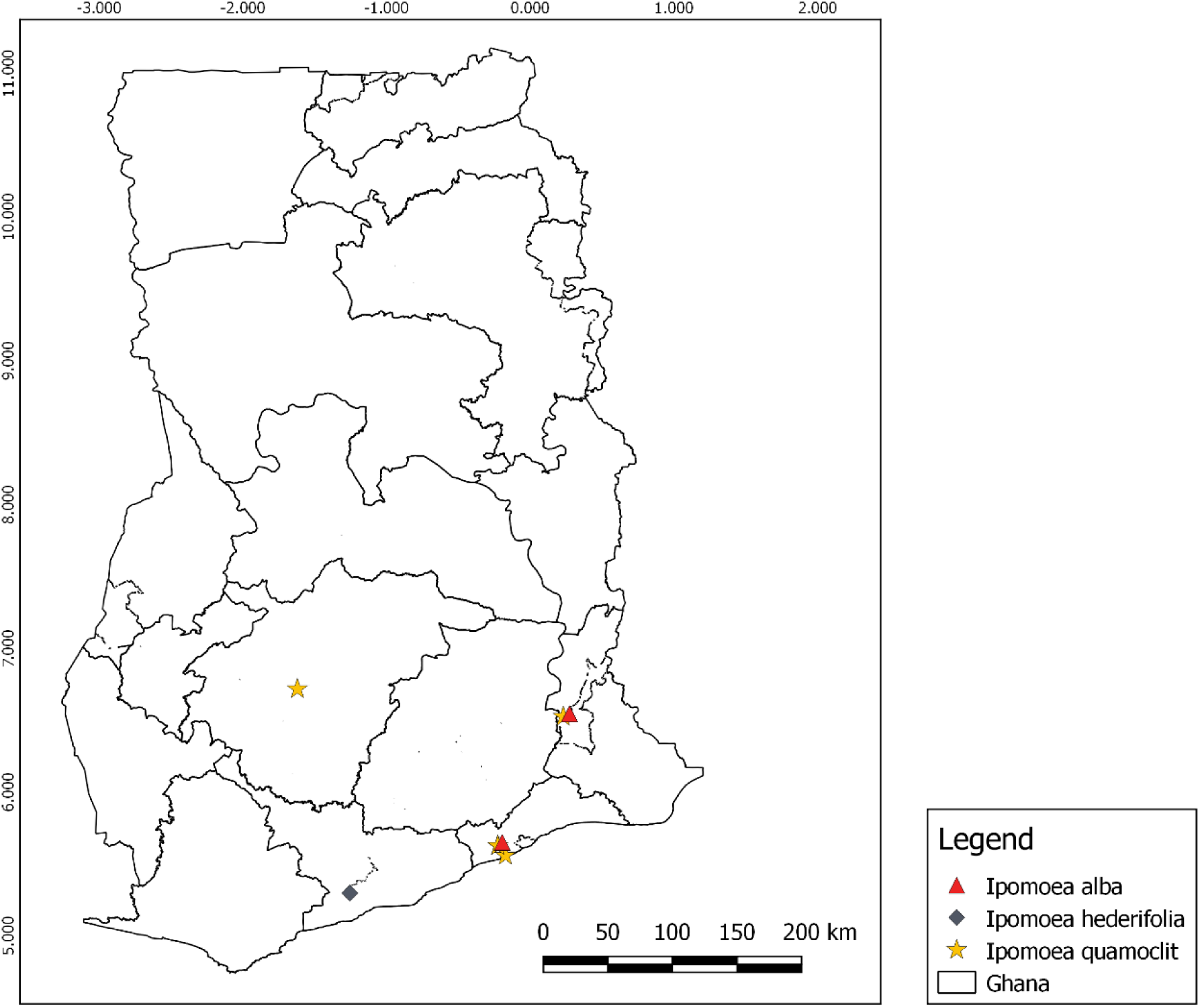
Distribution records of *I. alba*, *I. hederifolia* and *I. quamoclit* in Ghana.

#### HABITAT

Wild in secondary vegetation, but often cultivated for ornament; found in grassland, on riverbanks, along roadsides and in waste places.

#### VERNACULAR NAMES

*bona-nox* (Portuguese); *ndiami* (Nigeria: Efik); *bayugsns* (Sierra Leone: Kono); *bukbui* (Sierra Leone: Limba); *kpokpo-hina*, *hina: male* (Sierra Leone: Mende) (Burkill, 1985).

#### USES

The leaves of *Ipomoea alba* can be eaten as food, either cooked or uncooked, and probably as a supplementary famine food; whole herb is used to treat snake bite, the root bark is used as a purgative and the aerial part is used as an anti-pyretic, hypotensive and emollient; the leaves are used to treat headaches (Burkil, 1985).

#### CONSERVATION ASSESSMENT

LC: Least Concern (Canteiro, 2021).

#### PHENOLOGY

Flowers throughout the year, mostly January and February (Burkill, 1985).

#### SPECIMENS EXAMINED. GHANA

Greater Accra region: Legon, Legon hill, waste place, 1 Jan. 1956, *C.D. Adams* 3649 (GC); Krepi plains, 24 Jan. 1900, *W.H. Johnson* 548 (GC).

### 18. Ipomoea quamoclit L. (1753:159)

> Type: Netherlands (cultivated), Clifford s.n. (lectotype BM000558077!).

Annual *herb*. *Stems*: twining or prostrate, slender, angular, longitudinally ridged, glabrous. *Leaves* petiole 1 – 4 cm long, glabrous, with two leaf-like, subsessile, pseudo – stipules at the base; leaf pinnately compound, ovate to oblong in outline, 3.5 – 10 × 3 – 6 cm, glabrous, leaflets 8 – 19 per leaf, opposite to nearly so, linear to filiform, the two lower ones reflexed and bifid, glabrous. *Inflorescences* cyme, 1 – 3-flowered; peduncle slender, 3 – 8 cm long, glabrous; bracteoles deciduous, triangular, 1 – 3mm long. *Flower* pedicel 1 – 2.5 (– 4.2) cm long, slightly thickened at the apex, glabrous; sepals unequal, oblong to ovate-elliptic, apex obtuse and mucronate, glabrous, outer ones oblong, 4 – 5mm long, inner ones ovate – elliptic, slightly longer, c. 6mm long; corolla hypocrateriform, 2.5–5cm long, scarlet red-pink, glabrous, with a cylindrical tube, slightly and gradually narrowed towards the base, corolla lobes spreading, broadly triangular, acute and mucronate; stamens exserted, filaments of equal length, filiform, 19 – 23mm, widened and pubescent at the base, inserted 4 – 5 mm above the base of the tube; anther 1 – 2mm long, sagittate at the base; ovary ovoid, 1.5mm high, glabrous, 4-locular; style filiform, 19 – 30 mm long, glabrous, widened and articulated at the base. *Fruit* capsule, opening by 4 valves, ovoid to globose, 15mm x 18mm, with persistent style; seed fusiform, blackish, sparsely pubescent.

#### DISTRIBUTION

Widely distributed in the tropics and subtropics; native from Mexico to Central America. In Ghana: Greater Accra, Ashanti and Volta regions (Fig. 6). **New record for Ghana.**

#### HABITAT

A climbing annual, growing primarily in the seasonally dry tropical biome (POWO, 2024).

#### CONSERVATION STATUS (PRELIMINARY ASSESSMENT)

LC: Least Concern.

#### VERNACULAR NAMES

*pania oke* (Marquesas Islands); *batatilla* (Panama); *mayil manikkam* (Southern India); *ākāśamulla* (Malayalam); *tarulata*, *kamalata, kunjalata* (Bangladesh) (Prota4U.org).

#### USES

The leaves are used for general healing, treatment for haemorrhoids and for food, stems and roots for treatment of physical weakness, abnormal behaviour, sinking of voice, bleeding from cuts and wounds, piles, snakebites and constipation (Burkil, 1985).

#### SPECIMENS EXAMINED. GHANA

Greater Accra Region: Achimota Garden [5°37’23” N, 0°12’09”W], 2 Feb. 1931, *Asamany* 4 (GC); Achimota near Mr. Ateku’s bungalow [5°36’27”N, 0°14’03”W], 20 Jul. 1935, *G.K. Akpabla* 310 (GC); Ashanti Region: Near Wesley College Kumasi, [6°42’43”N, 1°37’21”W], 27 Apr. 1937, *Onyeama* 38 (GC); Volta region: Peki, 29 Mar. 1955, *E.D. Offori* 34 (GC).

### 19. Ipomoea hederifolia L. (1759: 925)

> Type: West Indies, Illustration of *Ipomoea foliis cordatis* in Plumier in Burman, Pl. Amer. 4: 82, t.93 f.2 (1756).

Annual *herb*. *Stems* twining, slender, ramified, slightly angular, glabrous or sparsely pilose. *Leaves* petiole 3 – 13 cm, glabrous to sparsely pubescent; leaves simple, sometimes 3-lobed, ovate to orbicular in outline, 2.2 – 8.3 cm x 2.1 – 8 cm, base cordate, apex acuminate and mucronate, margin entire, angular, coarsely dentate or deeply 3-lobed, glabrous to sparsely pubescent. *Inflorescences* axillary cymes, solitary to few-flowered; peduncle 1.3 – 16 (– 20) cm long, angular, glabrous to pubescent; bracteoles ovate, 1.5 – 2 mm, long mucronate. *Flower*: pedicel 0.3 – 1.2 (– 5) cm, enlarging in fruit; sepals slightly unequal, oblong-elliptic, 3 – 4mm long with a prominent awn, 2 – 4mm long, straight or slightly reflexed; corolla hypocrateriform, red scarlet, glabrous, limb 2 – 2.5 cm in diameter, shallowly 5-lobed, tube 2.8 – 4 cm long, narrowed below, slightly curved; stamens exserted, filaments c. 4 cm long, slightly widened at the base and covered with glandular hairs, anther 2mm long; ovary ovoid, glabrous, 2-locular, 4-celled, style c. 4.5cm long, stigmas 2-globose. *Fruit* capsule, opening y 4-valves, globose, 5 – 7mm in diameter, surrounded at the base by the persistent sepals; seeds trigonous, black, 3 – 4mm long, densely pubescent.

#### DISTRIBUTION

Native of tropical and subtropical America from the Southern United States to Argentina, now widely naturalized throughout the tropics. In Ghana: Central region (Fig. 6).

#### HABITAT

Found growing in waste places, thickets, cliffs, and locally established in riverine forest, roadsides, cultivated fields and disturbed areas.

#### VERNACULAR NAMES

*liseron hallier* (French), *trompetica roja* (Spanish), *fue kula* (Tongan), *amarra-amarra, corda-de-viola, batatarana, corriola* (Portuguese) (Burkill, 1985). **USES.** Roots are scraped and used as a remedy for stomach-ache; modified stems are also used to treat intestinal parasites (Rojas-Sandoval, 2016).

#### CONSERVATION STATUS (PRELIMINARY ASSESSMENT)

LC: Least Concern.

#### SPECIMENS EXAMINED. GHANA

Eastern region: Bunso, [6°16’47” N, 00°27’44” W], 17 Nov. 1995, *H.H. Schmidt* 1753 (GC); Eastern region: Agogo, 21 Dec 1928, *L.O*. *Deakin,* 8 (GC); Central Region: Assuantsi. 9 Feb 1928, *T.W. Williams* 1401 (GC).

### 20. *Ipomoea pyrophila* A.Chev. (1933: 235)

> Type: Mali, Koundian, *Chevalier 421* (P00434165!).

*Geophyte*, slender trailing herb; woody rootstock. *Stem* prostrate or climbing, densely covered in yellowish indumentum. *Leaves* petiole 1.7– 3 cm long, curved, densely pubescent with yellowish indumentum; leaf simple, oblong-ovate to broadly lanceolate, (1.4)2.5 – 5 cm x 1.2 – 2.2cm, apex retuse to obtuse and rounded, sometimes mucronate, base auriculate, sparsely yellowish pubescent above, trichomes appressed to the leaf surface, densely white and yellowish tomentose below, with hairs concentrated on the veins and midrib. *Inflorescences* axillary cyme, 1 – 3 – flowered; peduncle (1.1 –)1.3 – 5.2 cm, densely tomentose, with whitish spreading hairs; bracteoles narrowly elliptic to linear, up to 7 mm, pubescent. *Flower* pedicel 3 – 6 (– 13) mm long, slightly thicker than the peduncle, with similar pubescence; sepals subequal, lanceolate to oblanceolate, slightly narrower at the base, densely white tomentose, outer ones 6 – 10 (– 12) × 4 – 8 mm, inner ones 6×1.5 – 2 mm; corolla funnel-shaped, up to 2.8cm long, rosy-purple to pale violet with white lobes, sparsely pubescent on the outside; stamens included, filaments unequal, two longer ones of 9mm, three shorter ones of 7mm, gradually widening and weakly pubescent at the base; style included, 14mm long, glabrous; stigmas 2-globose. *Fruit* capsule, opening by 4 – valves, globose, 5 – 6mm in diameter, glabrous; seeds obloid, 3 mm long, pubescent.

#### DISTRIBUTION

West and Central Africa. In Ghana: Upper Eastern and Northern regions. New record for Ghana.

#### HABITAT

In savanna woodland after fires and in sub montane forest margins.

#### PRELIMINARY CONSERVATION ASSESSMENT

LC: Least Concern (proposed here).

#### SPECIMENS EXAMINED. GHANA

Northern region: Top of Damongo Scarp, Savannah woodland, 27 Mar.1953, *J.K. Morton* 8726 (GC); Upper Eastern region: On top of Gambaga hills, Savannah woodland, 5 Apr. 1953, *J.K*. *Morton* 8967 (GC).

### 21. *Ipomoea stenobasis* Brenan (1950: 228)

> Type: Uganda, Buganda Province, Mengo District, Entebbe, Nov. 1922, *Maitland 577* (holotype K).

Perennial *herb*, with a tap root. *Stems* prostrate, angular when young, becoming cylindrical, striate, fistulose, reddish, glabrous for the most part, puberulent at the insertion of the petiole. *Leaves*: petiole 2.5 – 3.6 cm long, glabrous; leaf simple, 5 – 7.5 (– 12) × 4.5 – 7.5 (– 11) cm, base cordate, apex acuminate and apiculate, margin entire, texture glabrous, covered with blackish dots on the lower surface, sparsely pubescent, especially along the veins, on both faces. *Inflorescences* axillary cyme, (1 –)3 – 5 flowered; peduncle 2 – 5 cm long; bracteoles deciduous, narrowly ovate-elliptic, 3 – 10mm long, glabrous, with hyaline margin. *Flower* pedicel 2 – 5 cm long, glabrous for the most part, base pubescent; sepals imbricate, unequal, elliptic, apex obtuse, outer ones shorter, 8 – 12 mm, with hyaline margin, inner ones longer, 12 – 15 mm; corolla magenta-pink, narrowly funnel-shaped, 3.5 – 6cm, entirely glabrous, base of the tube very narrow, with 2 – 2.5 mm of diameter; stamens included, filaments of unequal length; 11 – 16mm, widened and pubescent at the base; anther ovoid, 3mm long, sagittate at the base, obtuse at the apex; ovary 2 or 4-locular, glabrous; style 3 cm long, glabrous. *Fruit* capsule, 2.4-2.5cm x 1.1-1.6cm, surface striate, enclosed by the accrescent sepals, up to 22 mm long; seeds 3, 7-9mm long, ovoid, puberulent, especially around the hilum.

#### DISTRIBUTION

West Tropical Africa to Uganda (POWO, 2024). In Ghana: Brong Ahafo region (Fig. 4).

#### HABITAT AND ECOLOGY

Granite rocks in woodland and edges of swamps; 1,000– 1,200m.

#### CONSERVATION STATUS (PRELIMINARY ASSESSMENT)

LC: Least Concern.

#### SPECIMENS EXAMINED. GHANA

Brong Ahafo region, Nsemre F.R., between Borku and Anka, 20 Dec. 1954, *C.D. Adams* 2843 (GC).

### 22. *Ipomoea violacea* L. (1753: 161)

> Type: Icon in Plumier Pl. XXXIV (lectotype: Bibl. Univ. Groningen).

Perennial, glabrous, *herb*. *Stems* woody at the base, twining or prostrate, often longitudinally wrinkled, but otherwise smooth. *Leaves* petiole glabrous, 3.2 – 11 cm; leaf simple, cordate, circular or ovate, 5–16 × 5–14 cm, apex acuminate, mucronate, rounded or rarely angular, base cordate, glabrous. *Inflorescences* 1 – to few flowered; peduncle 2.5 – 4.5(−7) cm; sepals subcircular, equal or outer two shorter, 1.5 – 2.5 cm, margins hyaline, coriaceous, apex obtuse or emarginate, mucronulate. *Flower* pedicel 1.5-5 cm, thickened in fruit; corolla night flowering, salver-shaped or very narrowly funnel-shaped, white or pale greenish-yellow, with green midpetaline bands, 9-12 cm long, glabrous; ovary glabrous, stigma 2 –globose; filaments inserted near the base of the corolla tube. *Fruit* capsule, opening by 4-valves, ovoid to globose, glabrous, pale brown, 2 – 2.5 cm; seeds sub-trigonous, 1 – 1.2 cm long, black, densely short tomentose, edges with ca. 3 mm long sericeous hairs.

#### DISTRIBUTION

Native in tropical and subtropical coasts. In Ghana: Western region (Fig. 7).

**Figure 7.**
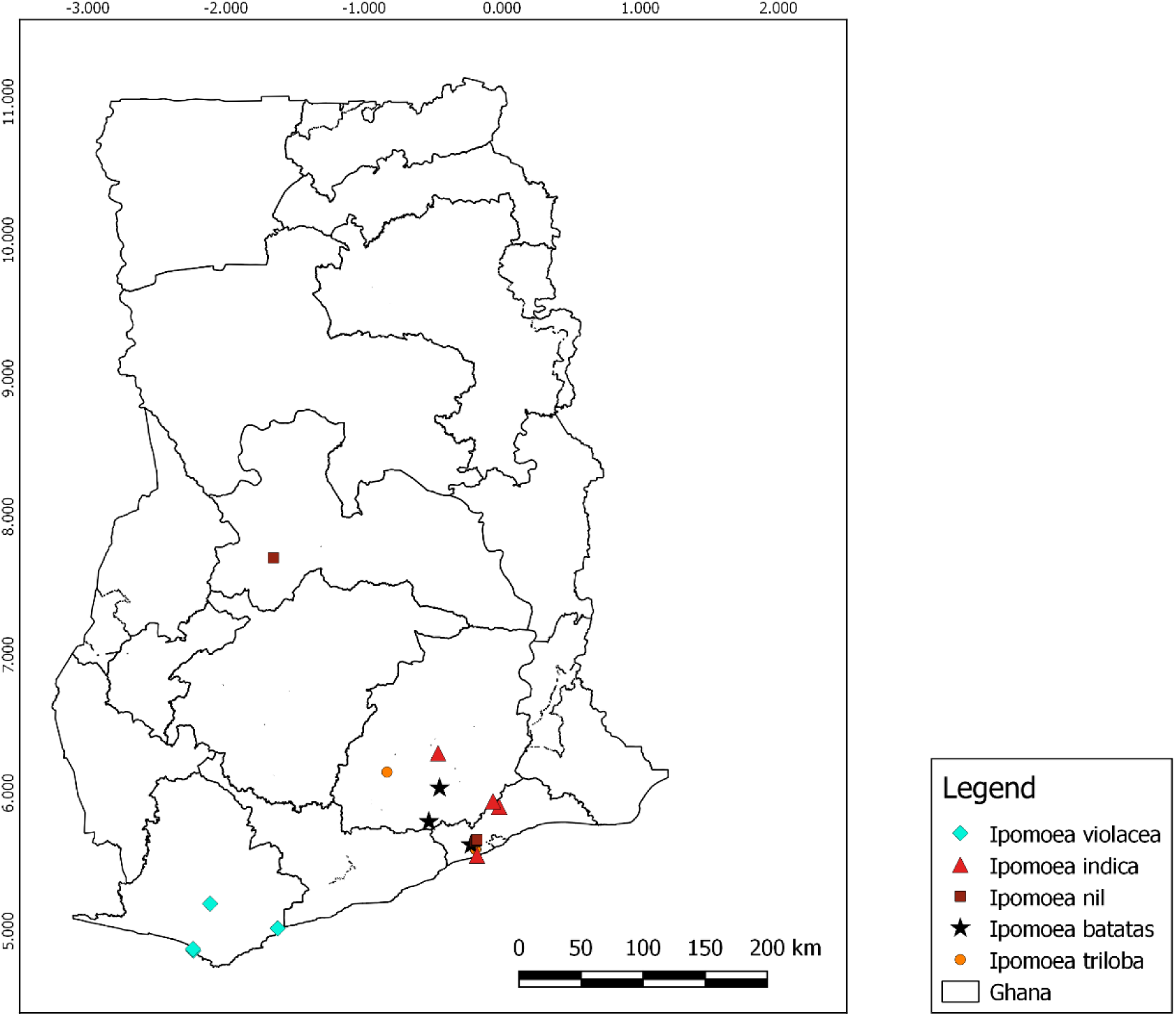
Distribution records of *I. violacea*, *I. indica*, *I. nil*, *I. batatas* and *I. triloba* in Ghana.

#### VERNACULAR NAMES

beach moonflower, morning glory, sea moonflower (English) (Michaels, 2022).

#### USES

The seed contains small quantities of the hallucinogen LSD, this has been used medicinally in the treatment of various mental disorders; the leaves and the tuber serve as food (Michaels, 2022).

#### HABITAT

Coastal bushland; beaches, seaside thickets, edges of brackish rivers and lagoons; near sea level to 100m (Heine, 1963).

#### CONSERVATION STATUS (PRELIMINARY ASSESSMENT)

LC: Least Concern.

#### SPECIMENS EXAMINED. GHANA

Western region: Axim, strand vegetation near beach, 1 Mar.1934, *F.R. Irvine* 2565 (GC); *loc. cit*., 1 Apr. 1965, *J.K. Morton* A2092 (GC); *loc. cit*., 19 May 1956, *J.K. Morton* 2092 (GC); Shama, 15 May 1965, *D. Hall* 2982 (GC).

### 23. *Ipomoea obscura* (L.) Ker Gawl.(1817: t. 239)

> Type: Icon. Dillenius in Hort. Eltham.: tab. 83, fig. 95 (1732).

Perennial *herb*. *Stems* prostrate to twining, filiform, cylindrical, nearly woody at base, pilose or glabrescent. *Leaves* petiole 1-11 cm long, slender, pubescent or glabrescent; leaf entire or slightly undulate, ovate, rarely linear-oblong, 2.5-8.5 cm x 0.4-7.5 cm, cordate with rounded auricles at the base, acuminate or apiculate and mucronate at the apex, margin often ciliate, membranous, pubescent or glabrescent on both surfaces. *Inflorescences* axillary cymes, 1-several-flowered, pubescent, 1-4(−5.5) cm long; peduncle 1-8 cm long, slender, glabrous, pubescent or thinly pilose; bracteoles triangular, acute, 1-2 mm long. *Flower* pedicel 1-2 cm long, at first erect but in fruit relaxed and thickened towards the apex, sometimes minutely verrucose, glabrous or less commonly, pubescent or pilose; sepals subequal, ovate, ovate-orbicular, ovate-lanceolate or lanceolate, apex acute or apiculate, 4-8 mm long, 1.7-4 mm long, often wrinkled or muricate, margin scarious, glabrous or pilose with long white trichomes, in fruit all somewhat accrescent, ultimately often spreading or reflexed, two outer sepals shorter, ovate, apex acute to shortly acuminate or mucronate, inner sepals ovate-elliptic, apex obtuse, occasionally mucronate; corolla funnel-shaped, 1.5-2.5 cm long, yellow, orange, cream or white, with or without a dark purple centre, often weakly lobed, 3-4 cm in diameter, midpetaline bands sparsely pubescent at the apex; stamens included, filaments unequal, two longer, three shorter, widened and pubescent at the base, anthers ovoid, base sagittate, 2-3 mm long; ovary elongated to rounded, distinctly prolonged at apex, 2.8-5 mm long, glabrous, 2-locular, 4-ovuled; style filiform, 7-8 mm long, glabrous, stigmas 2-globose. *Fruit* capsule globose-ovoid, 7-12 x 5-10 mm, glabrous, crowned by persistent style base; s*eeds* ovoid, 4-5.5 mm long, black, appressed pubescent.

#### DISTRIBUTION

Native in tropical and subtropical regions of Africa and Asia. In Ghana: Western, Central and Greater Accra regions (Fig. 8).

**Figure 8.**
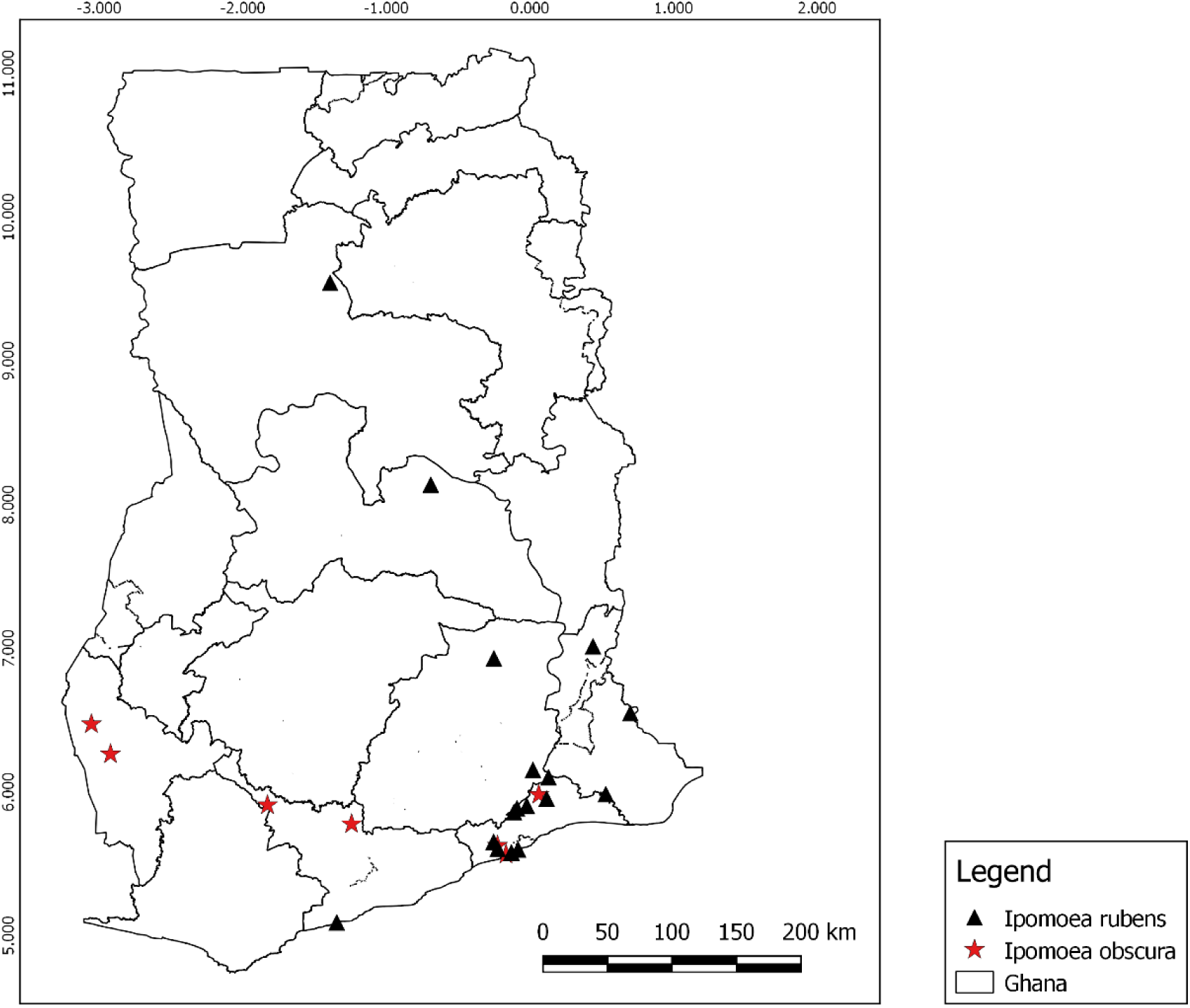
Distribution records of *I. rubens* and *I. obscura* in Ghana.

#### HABITAT

In forests, shrub lands, and rocky areas e.g., inland cliffs or mountain peaks at 500-2,200m altitude (Demissew, 2006; Shimpale *et al.,* 2012).

#### CONSERVATION STATUS

LC: Least Concern (Allen, 2017).

#### VERNACULAR NAMES

*ododo oko, ododo owuro* (Nigeria: Yoruba); *Ògbànanì* (Nigeria: Igbo) (Burkill, 1985).

**USES.** The leaves are eaten as food (Burkil, 1985); it is also a widely cultivated ornamental (Allen, 2017).

#### SPECIMENS EXAMINED. GHANA

Greater Accra region: between Achimota and Little Legon, 14 Oct. 1955, *C.D*. *Adams* 3324 (GC); Legon village, 1 Mar. 1931, *F.R. Irvine* 1598 (GC); Ayikuma to Shai Hills Road, 4 Dec. 1952, *J.K*. *Morton* 8069 (GC); on top of Shai Hills, 8 Jan. 1953, *J.K. Morton* 8274 (GC); Upper West region: Wa South, 25 Jun. 1950, *C.D*. *Adams* 794 (GC).

### 24. *Ipomoea rubens* Choisy(1833: 463)

> Type: India, Silhet and Goalpara, *Wallich 1421* (lectotype G, isolectotypes K, W, G).

Perennial *herb*. *Stems* rather woody, finely striate when dry, terete, 1-3 mm in diameter, up to 4 m long, densely short-pilose with soft greyish hairs. *Leaves* petiole 3-7 cm long, slender, pilose like the stems; leaf entire or shallowly 3-lobed, broadly ovate to circular, 5-15cm x 4-12 cm, cordate with rounded auricles at the base, apex acuminate and mucronate, upper surface pubescent to densely villose, lower surface less densely pubescent to glabrous, with greyish hairs; 7-9 pairs of secondary veins. *Inflorescences* compact, axillary, sub-umbelliform, cymes, 1-few-flowered; peduncle 2-15 cm long, cylindrical, pilose as the stems; bracteoles triangular, 3-7 mm long, pubescent, caducous. *Flower* pedicel 5-17 mm long, pilose; sepals subequal, ovate, apex obtuse or minutely mucronate, 6-8(−11) x 3-6(−10) mm, pilose at the apex but not ciliate, persistent in fruit, outer sepals accrescent to 16 mm in fruit, ovate-deltoid, apex acute, inner sepals apex obtuse, margin distinctly hyaline; corolla funnel-shaped, 4-5 cm long, purple or mauve with dark purple centre, limb 4-5 cm in diameter, sparsely pilose, midpetaline bands with silky long trichomes; stamens included, filaments unequal, the longer ones 8-9 mm long, shorter ones c. 3 mm long, widened and pubescent at the base; anthers obloid, c. 2 mm long; base sagittate; ovary glabrous, 2-locular, 4-ovuled; style filiform, 15-17 mm long, stigmas 2-globose. *Fruit* capsule, opening by 4 valves, globose, 12-13 mm in diameter, glabrous; s*eeds* ovoid, 6-9 mm, densely pilose, with white or yellowish hairs c. 2.5 mm long.

#### DISTRIBUTION

Native to tropical and subtropical regions of Africa and Asia. In Ghana: Brong Ahafo, Greater Accra, Northern, Volta and Central regions (Fig. 8).

#### CONSERVATION STATUS

LC: Least Concern (Ghogue, 2020).

#### PHENOLOGY

Flowering throughout the year.

#### HABITAT

*Ipomoea rubens* occurs in humid, sub-humid, and dry freshwater wetlands (Demissew, 2006).

#### ETYMOLOGY

“*rubens”* means red-coloured; the specific epithet refers to its red/purple corolla (Wamucii, 2024).

#### VERNACULAR NAMES

*otjiherero*, *otjinatjoruhona* (Namibia) (Gonçalves, 1987).

#### USES

The leaves, shoots, and leafy stems are eaten by primates such as chimpanzees (Calabuig, 2014).

#### SPECIMENS EXAMINED. GHANA

Northern Region: White Volta at Daboya Tamale, 16 Jan. 1996, *A.A. Ente* & *C.W*. *Agyakwa* VBS 419 (GC); Greater Accra region: Accra plains, little lake Volta [6°09’48’’N, 0°03’54’’E], alt. 50m, 12 Nov. 1994, *C.C.H*. *Jongkind*, *D*. *Abbiw* & *C.M*. *Markwei* 1855 (GC); Brong Ahafo region: Yeji, 11 Apr. 1964, *J.B. Hall* 1299 (GC); *loc. cit*.,13 Aug. 1963, *B.K. Adom Boafo & M. Ansa-Emmim*, VBS 191 (GC); *loc. cit*., 19 May 1965, *A.A*. *Ente* & *C.W. Agyakwa* VBS 2115 (GC).

### 25. *Ipomoea triloba* L. (1753: 161)

> Type: Icon in Sloane, Voy. Jamaica 1: t. 97, f. 1 (1707).

Annual *herb*. *Stem* prostate, glabrous or puberulous, especially on the nodes. *Leaves* petiole angular, 2.5-6 cm long, glabrous, sometimes tuberculate; leaf simple, broadly ovate to circular in outline, apex attenuate, base cordate, margin coarsely dentate to deeply 3-lobed, 2.4-7.5 x 3-6.4 cm, glabrous or sparsely pubescent. *Inflorescences* axillary, umbelliform, cyme, few-to many-flowered; peduncle (1-) 2.5-5.5 cm long, angular, verruculose at the apex, glabrous; bracteoles lanceolate to oblong-ovate, 1–2 mm long, glabrous. *Flower* pedicel 5–8 mm long, sometimes angular, verruculose, pubescent at the base; sepals subequal, 5-8 mm, outer sepals oblong, slightly shorter, inner sepals elliptic-oblong, all glabrous or sparsely pilose outside, margins fimbriate, apex obtuse or acute, mucronulate; corolla funnel-shaped, 1.5-2 cm, pink or mauve, with dark purple center, glabrous, limb obtusely 5-pointed; stamens included; ovary pubescent. *Fruit* capsule, opening by 4-valves, spherical to ovoid, globose, 3–10 mm long, 5– 9 mm wide; seeds spherical, rounded, 2-3.5 mm long, dark brown, glabrous.

#### DISTRIBUTION

Native from Mexico to Brazil, and Caribbean (Alencar *et al.,* 2021; POWO, 2024). In Ghana: Greater Accra and Eastern regions (Fig. 7). **New record for Ghana.**

#### HABITAT

The species is found in grasslands, upland cultivated fields, roadsides, waste lands and disturbed sites; it is a climbing annual or perennial and grows primarily in the seasonally dry tropical biome (CABI, 2021).

#### CONSERVATION ASSESSMENT

LC: Least Concern (Contreras & Wood, 2019).

#### USES

The whole plant is used as painkillers (Burkill, 1985).

#### SPECIMENS EXAMINED. GHANA

Eastern region: Kade Agricultural Research Station [6 °8’28”N, 00°53’56”W], alt. 200m, 22 Dec. 1996, *H.H. Schimdt, M. Merello, J. Amponsah & M. Chintoh* 2271 (MO, GC); Greater Accra, Maamobi, 4 May 1976, *J.B. Hall* 43692 (GC).

### 26. *Ipomoea batatas* (L.) Lam. (1793: 465)

> Type: Inde, Herb. Linn. LINN 77.5 (S-LINN).

Perennial *herb,* with underground, fusiform to ellipsoid, yellow or reddish edible storage roots, colour depending on cultivar. *Stems* prostrate, ascending or rarely twining, angular when young, 1-2mm in diameter, often rooting at the lower nodes, striate, glabrous or glabrescent. *Leaves* petiole 4-20(−50) cm long, glabrous or puberulous; leaf simple, triangular to broadly ovate in outline, entire to 3-5-lobed, palmatifid or palmatisect, 4-14 x 4-16cm, apex acute to acuminate and mucronate, base truncate or cordate, lobes triangular to lanceolate, glabrous to slightly pubescent on both surfaces. *Inflorescence* axillary cymes, 1-several-flowered; peduncle 3-18cm long, stout, angular, glabrous or pubescent; bracteoles narrow, oblong, acute, 2-3mm long, early deciduous. *Flower* pedicel 3-12 mm long; sepals subequal, oblong to elliptical-oblong, apex acute and distinctly mucronate, 7-12mm long, 3-5mm wide, subcoriaceous, glabrous or pilose on the back and fimbriate, persistent in fruit, outer sepals oblong to elliptic-oblong, 7-12 x 2-3mm, inner sepals elliptic-oblong or ovate oblong, 9-12mm long, 4-5 mm wide; corolla funnel-shaped, 3-5cm long, violet or lilac to white, with dark purple center, glabrous; stamens included, filaments unequal, two longest 8–14mm long, three shortest 6– 9mm long, widened and pubescent at the base, anthers obloid, 2-3mm long, base sagittate; ovary ovoid, 1.5-2mm long, sparsely pubescent, with long spreading hairs, 4-celled; style filiform, 15-20mm long, glabrous, stigmas 2-globose. *Fruit* capsule ovoid, opening by 4-valves, 8-12mm long; s*eeds* ovoid to irregularly trigonous, 4-7mm long, black, glabrous.

#### DISTRIBUTION

Native from Mexico to Venezuela and Ecuador (POWO, 2024). In Ghana: Greater Accra and Eastern regions (Fig. 7).

#### HABITAT AND ECOLOGY

Sweet potatoes are grown at low to medium elevations of up to 1,800 m in regions with moderate rainfall or in damp environments. Beyond this altitude, to 2,200 m and higher, it is grown as a fodder crop. Though its precise origin is unknown, this plant is widely cultivated across the tropics. When plants are not cultivated, they are typically found on roadsides close to homes and farms, as well as in abandoned fields (Cartabiano-Leite *et al.,* 2020).

#### CONSERVATION STATUS

DD: Data deficient (Rowe *et al.,* 2019).

#### VERNACULAR NAMES

*sweet potato* (English), *patate douce* (French), *lémongho, futa* (Mindumu), *lifita* (Bavili), *imongo* (Banzabi), *égwèta* (Ivéa, Mitsogo), *lungu* (Bapunu) (Prota4U.org).

#### USES

The edible roots are roasted, fried, or boiled before consumption; they are processed to make crisps and sweet potato fries (chips); additionally, they provide starch, which is used in Korea to produce *dang myun* noodles; in New Guinea and Southeast Asia, sweet potato leaves are eaten as a vegetable; owing to their lovely flowers and foliage, certain sweet potato cultivars are planted as ornamentals (Carvalho *et al.,* 2023).

#### SPECIMENS EXAMINED: GHANA

Greater Accra region: Maamobi, 4 May 1976, *J.B. Hall* 43692 (GC); Volta region: Kade agricultural research station [6°08’28’’N, 00°53’56’’W], alt. 200m, 2 Dec. 1996, *H. Schmidt, J. Amponsah, M. Chinto* 2271 (GC).

### 27. *Ipomoea nil* (L.) Roth (1797: 36)

> Type: Icon. Dillenius, Hort. Eltham. 96, t. 80, f. 91 (1732).

Annual *herb*. *Stems* twining, terete, 1 mm in diameter, pubescent with bristly simple whitish hairs. *Leaf*: petiole 1.5-4 cm long, densely villose with bristly simple whitish hairs; leaf simple, entire or 3-lobed, ovate to circular in outline, 4-14 x 3-13.5 cm, apex acuminate, base cordate, pubescent with appressed whitish simple hairs on both surfaces, denser below. *Inflorescence* axillary lax cyme, 1-few-flowered; peduncle 2-10 cm long, densely hirsute with whitish hairs; bracteoles linear to filiform, 5-8 mm long. *Flower* pedicel 5-10 mm long, with similar pubescence to the peduncle; sepals subequal, linear-lanceolate, 15-28 x 3.5 mm wide, densely pilose with spreading hairs, persistent in fruit; corolla funnel-shaped, (4-)5-6 cm long, blue to mauve with paler tube, often white inside, glabrous; stamens included, filaments unequal, the longest 20-22 mm long, the shortest 12-15 mm long, widened and pubescent with long hairs at the base; anthers obloid, base sagittate, 3 mm long; ovary ovoid, 3-locular, glabrous; style filiform, stigmas 2-globose. *Fruit* capsule, opening by 3 valves, ovoid to globose, 8-12 mm long, glabrous, surmounted by the persisting base of the style, enclosed by the calyx. *Seed* obovoid-trigonal, 4.5-6 mm long, black, puberulous with fine greyish hairs.

#### DISTRIBUTION

Native in tropical and subtropical America (POWO, 2024), cultivated as ornamental, or escaped from cultivation, elsewhere. In Ghana: Greater Accra and Brong Ahafo regions (Fig. 7).

#### HABITAT AND ECOLOGY

A climbing annual, growing primarily in the seasonally dry tropical biome (POWO, 2024).

#### CONSERVATION STATUS (PRELIMINARY ASSESSMENT)

LC: Least Concern.

#### USES

The seeds of *Ipomoea nil* are used as laxatives and purgatives (Burkil, 1985). In Indonesia, *Ipomoea nil* has environmental and social uses, as animal food, a poison and a medicine and for food. (POWO, 2024).

#### SPECIMENS EXAMINED. GHANA

Greater Accra region: La Nkwantanang Madina, Legon Botanical Gardens, 19 Dec. 1969, *A.A. Enti*, 42762(GC); *loc. cit.*, 25 Oct. 1961, *J.K. Morton* 5010 (GC); *loc. cit*., 16 Jul. 1965, *M. Abedi Lartey* MAL/1(GC); Ashanti region: Bobiri F.R., 24 Aug. 1963, *A.A. Enti* 35246 (GC).

### 28. *Ipomoea indica* (Burm.) Merr. (1917: 445)

> Type: Besler, Aest. Ord. 13, fol. 8. II (1613).

Perennial *herb*. S*tems* twining or prostrate, 1-1.5mm in diameter, rooting at the nodes, pubescent with retrorse hairs, subligneous at the base. *Leav*es: petiole 2-10cm long, pubescent; lamina simple, ovate in outline, entire or 3-lobed, 5-12 x 3-15cm, cordate at the base, apex acuminate, pilose to glabrescent on both surfaces, more densely pubescent below. *Inflorescences* dense axillary cymes, few-many-flowered; peduncle 0.5-15cm long; bracteoles linear to lanceolate, or ovate-lanceolate. *Flower* pedicel 2-10cm long, pubescent like the stem; sepals subequal, lanceolate, apex caudate, acuminate, 1.5-2.3cm long, herbaceous, glabrescent, persistent in fruit, outer sepals ovate, apex long acuminate, 9–12mm long, pubescent, inner sepals narrower; corolla funnel-shaped, 5-8cm long, blue or mauve-purple, often red-tinged, tube whitish at the base, limb flaring, glabrous, lobes broadly rounded, notched at apex; stamens 5, included, filaments unequal, 17–30mm, broadened and hairy at the base, anthers obloid, 3.5– 5(–5.3)mm long; pollen spinulose, pantoporate; *disc* annular, lobed; ovary ovoid, 3-locular, 1-1.5mm long, glabrous; style filiform, 30-33mm long, glabrous, stigmas 2-globose. *Fruit* capsule ovoid-globose, opening by 3 valves, 8-10mm in diameter, glabrous; s*eeds* 6, ovoid to ellipsoidal, 4-6mm long, brown-black, covered with an appressed pubescence.

#### DISTRIBUTION

Native to tropical and subtropical America, cultivated as ornamental or escaped from cultivation elsewhere. In Ghana: Greater Accra and Eastern regions (Fig. 7).

#### HABITAT AND ECOLOGY

Found in roadside thicket sand along the borders of moist woodlands.

#### CONSERVATION STATUS (PRELIMINARY ASSESSMENT)

LC: Least Concern.

#### USES

Cultivated as ornamental, sometimes escaped from cultivation or sub-spontaneous (Mwanga *et al.,* 2022).

#### VERNACULAR NAMES

blue dawn morning-glory, *ocean-blue morning-glory* (English) (Mwanga *et al.,* 2022).

#### PHENOLOGY

It flowers in most months of the year (Wood *et al.,* 2015).

#### SPECIMENS EXAMINED: GHANA

Greater Accra region, Accra Plains, 15 Feb. 1954, *G.K. Aghemaheie* (GC); Christiansborg, 17 Apr. 1954, *F.W. Engmann* (GC); Eastern region: Koforidua [6 ° 16’47”N, 00 ° 27’44”W], alt. 150m, 17 Nov. 1995, *H. H. Schmidt, J. Amponsah & A. Welsing* 1751 (MO, GC); Larteh, 18 Oct. 1900, *W. H. Johnson* 822 (GC).

## Acknowledgements

Authors are grateful to the Schroder Foundation and Mallinckrodt Foundation for fully funding the “Plant Taxonomy Skills for Ecology and Conservation 2022” course, co-organised by Royal Botanic Gardens, Kew and the University of Ghana. G. Ameka, R. Borosova and A.R.G. Simões thank the International Association for Plant Taxonomy (IAPT) for the “Biodiversity Challenge” 2023 Award which allowed essential improvements to the GC Herbarium, and motivated the Botany students at the University of Ghana to continue developing taxonomic work, such as the present study. Thanks also due to Rafael Felipe de Almeida (Royal Botanic Gardens, Kew) for useful comments on the manuscript.

## Notes

### Competing Interest Statement

The authors have declared no competing interest.

